# Morphological profiling of all human and mouse miRNAs in 24M cells

**DOI:** 10.1101/2025.11.14.687149

**Authors:** Jesko Wagner, Claire L. Smillie, Tamara M. Sirey, Ava Khamseh, Chris P. Ponting, Sjoerd V. Beentjes

## Abstract

MicroRNAs (miRNAs) are post-transcriptional regulators of gene expression whose contributions to cell biology remain underexplored. Here, we used Cell Painting to quantify the morphological effect of 2,565 human and 1,900 mouse miRNA mimics on 24 million cells across five cell lines. To do so, we developed a novel single-cell morphological profiling analysis framework, involving stringent batch correction, feature selection, and hit calling. With this, we discovered that 9% of human and 15% of mouse miRNA mimics significantly alter cell morphology in at least one cell line. Eighteen miRNAs caused significant changes in multiple cell lines, including eight orthologous miRNAs that altered morphology in both human and mouse cells. Among the replicating miRNAs were human and mouse miR-155-5p, which affected morphology in at least one replicate of four cell lines. As expected, miRNAs with identical seed sequences induced more similar morphological changes than miRNAs with different seeds. Also, morphological changes were associated with cytotoxicity and annotation confidence. This comprehensive single-cell morphological resource will help elucidate the cellular functions of human and mouse miRNAs.

## Introduction

MicroRNAs (miRNAs) are small (∼22 nucleotide) non-coding RNAs that are important modulators of post-transcriptional regulation^1^. Bound to an Argonaute family member, they form the RNA-induced silencing complex (RISC) that binds target mRNAs through partial sequence complementarity to the miRNA seed region (nucleotides 2–8)^2,3^. Canonically, binding triggers degradation of the target mRNA or inhibits its translation, thereby reducing the abundance of the protein encoded by the bound mRNA^4^. MiRNA-mediated post-transcriptional regulation may be widespread: approximately 60% of all human protein-coding transcripts contain a conserved miRNA binding site^5^. Many of the 2,654 human miRNAs discovered to date^6^ play roles in cell proliferation^7^, differentiation^1^, signaling^8,9^, metabolism^10,11^, embryonic development^12^, and disease^13,14^. However, the effect of most miRNAs is difficult to study; many miRNAs affect proliferation and survival^15^, but their exact cellular role often remains unclear.

Associating miRNAs to phenotypes presents three challenges. First, although most miRNAs affect the abundance of individual mRNAs only modestly, they could substantially alter phenotypes by co-ordinately modulating the abundance of hundreds of mRNA targets^14,16,17^. Second, the miRNA literature is contaminated by paper mill publications and by loci incorrectly annotated as miRNAs^2,17–21^. Third, forward genetic studies of miRNA function are potentially hampered by cell-type specificity of miRNA action^21^. To interrogate miRNA function beyond their impact on cell survival and gene expression, comprehensive data resources and methods are required to link miRNAs to phenotypes of diverse cell types.

Cell Painting is a morphological profiling assay based on high-content microscopy that can compare cellular morphologies across thousands of perturbations. It uses six fluorescent dyes to stain eight cellular components in five channels, which yield numerical morphological descriptors (“profiles”) for each cell^22,23^. These profiles capture cell size and shape information as well as fluorescent intensities of cell organelles and compartments, which permit detection of changes in cell morphology. Cell Painting can scale to millions of cells, which could systematically reveal miRNA function. In a smaller experiment, Singh et al.^24^ used Cell Painting and 315 short hairpin RNAs targeting 41 genes to show that seed sequence and off-target effects determine cell morphology. Insights gained from morphological profiling are, however, usually limited due to cell profiles being averaged per well or treatment, rather than cells being analysed individually. Averaging could mask subtle changes occurring in subpopulations of cells. This is particularly relevant to experiments using transfection whose efficiency is imperfect^25^. Retention of single-cell information throughout an analysis can improve analysis accuracy^26,27^, motivating a single-cell morphological analysis of miRNA activity.

Here, we used single-cell analysis with Cell Painting to test the effects of 2,565 human and 1,900 miRNA mimics in three human and two mouse cell lines. We found that 226 (9%) human and 286 (15%) mouse miRNAs induce significant changes in cell morphology in at least one cell line, including 18 miRNAs that induce changes in two cell lines. Seed sequences often dictated morphological changes, as seen for five paralogs of hsa-miR-518. This is the first single-cell study to comprehensively associate miRNAs with cellular morphology. It also highlights the steps needed to analyse single-cell data from complex morphological profiling experiments.

## Results

We measured the effects on single-cell morphology from transfecting each of 2,565 miRNA mimics in three human cell lines, and 1,900 miRNA mimics in two mouse cell lines (Figure 1A). We set up 384-well plates such that each miRNA was administered four times per cell line: in two wells of one plate, and in two wells of a second plate seeded with a different passage of cells (termed “replicates” throughout), yielding a total of 134 plates (Figure 1B). We assessed effects of individual miRNA mimics (hereafter miRNAs) on cell morphology using the Cell Painting assay. This is a high-content image-based profiling assay that stains eight components of the cell – DNA, endoplasmic reticulum (ER), cytoplasmic RNA, nucleoli, actin, Golgi apparatus, plasma membrane and mitochondria – and is imaged in five channels (Figure 1C). Then, we obtained 205,824 fluorescent microscopy images of 24,127,517 cells. From these, we extracted 1,386 descriptors of cell morphology (“features”), such as intensity of stains and cell size, which together form the morphological profile of each cell (Figure 1D).

**Figure 1.**
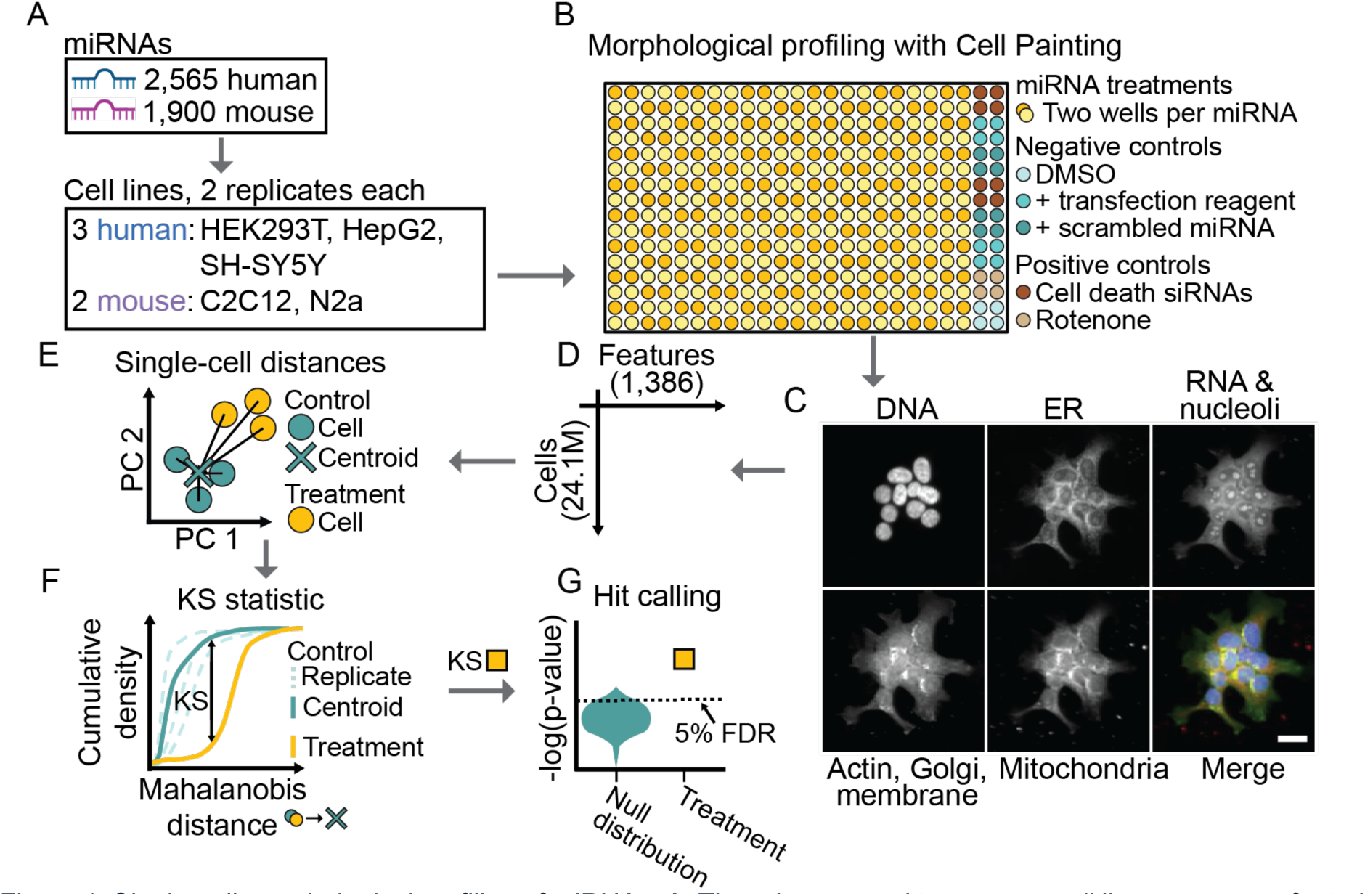
Single-cell morphological profiling of miRNAs. A: Three human and two mouse cell lines were transfected with 2,565 human or 1,900 mouse miRNA mimics, respectively. Two experimental replicates per cell line were used. B: Each miRNA was applied to two wells, requiring 134 plates in total. C: After Cell Painting, eight cellular compartments were visualised with fluorescent microscopy, as exemplified with images of HEK293T cells. The scale bar is 25 μM. D: Single-cell profiles were computed for 24.1M cells across 1,386 features. E: After image QC and batch correction (Methods), a Mahalanobis distance for each single cell was calculated from its location on the first 10 principal components (PC) to the average location of negative control cells (control centroid). These distances were calculated in a space of PCs derived from 1,386 features. F: The Kolmogorov-Smirnov (KS) statistic was used to compare the treated cells’ phenotypic distance from the control centroid to the distances of negative control cells. G: To estimate the significance of these phenotypic shifts (“hit calling”), a background distribution was built from pairwise KS tests between negative control wells across all plates of a replicate experiment. Treatments were considered significantly different from negative controls when their p-value was below the 5% FDR threshold determined by the background distribution.

### Single-cell morphological profiling analysis framework

To understand which miRNA treatments caused quantifiable changes in cell morphology, we needed to overcome three challenges. First, images had to be quality controlled to remove artifacts like dust precipitation, which we addressed with a nearest-neighbour approach that identified and removed affected images (1-7%, depending on cell line, Supplementary Figure S1). Second, plate-positional and batch effects caused by technical factors such as photobleaching, common for Cell Painting experiments^25^, were addressed. Left unaccounted for, these technical effects can overshadow treatment effects^28^. We performed batch correction using a linear model and selected features that were not significantly associated with plate position or batch (Methods and Supplementary Figure S1). Third, we established methods that determine whether a miRNA treatment significantly altered cell morphology in successfully transfected cells.

To do so, we developed scmorph, a Python package for the single-cell analysis of morphological profiles^29^. This measures morphological heterogeneity in two cell populations: (i) cells treated with a scramble miRNA, termed “negative controls”, and (ii) cells treated with a miRNA of interest. More specifically, we first reduce the dimensionality of the morphological profiles using principal component analysis (PCA), before computing the Mahalanobis distance of each cell to the average location of negative control cells on the first 10 PCs (“centroid”; Figure 1E). This yields a vector of distances for each treatment (“treatment distances”) and the negative control (“control distances”). For each treatment we then summarise its difference to negative controls by performing the Kolmogorov–Smirnov (KS) test, comparing the treatment and control distances (Figure 1F). This test measures the probability of the null hypothesis that treatment and control distances are drawn from the same distribution, i.e. that there are no differences between treated and negative control cells.

We then tested whether a miRNA affected cell morphology (i.e., was a “hit”) by combining cells from two technical controls (on the same plate) and comparing it to a negative control, which required new methods to overcome the high sensitivity of morphological profiling to minor technical variation^25^. First, we decided which of three negative controls to use as a reference: DMSO only, DMSO with transfection reagent (DMSO+TR), or DMSO+TR with an additional scrambled miRNA. The three negative controls yielded similar results when compared to positive controls (Supplementary Figure S2). miRNAs could significantly change morphology differently across the negative controls (Supplementary Figure S3). However, 96% of miRNAs yielded the same result (hit or no hit) using each of two biologically relevant negative controls. We opted to report results from wells treated with scrambled miRNAs as negative control to most closely mimic treatment conditions.

Hit calling was performed using an empirically controlled false discovery rate (FDR). This involved building a null distribution of pairwise tests of negative control wells, between which we expect no morphological differences (see Methods). If a miRNA’s p-value was smaller than the smallest 5% of p-values of the null distribution then it was considered significant (FDR < 5%, Figure 1G). Lastly, we defined miRNAs as hits only when they were significant in both replicates of a cell line, which reduces false positives (Methods).

### Cell death siRNAs impact morphology of surviving cells

To test our single-cell analysis methods, we first investigated whether treatment of HEK293T cells with a cocktail of cell death-inducing siRNAs results in significant changes to cell counts and cell morphology. This positive control treatment was applied to specified wells on every plate; the treatment also served as a transfection control. As expected, 24 h after exposure to the siRNA cocktail the average cell count was significantly reduced by 39% relative to a negative control of scrambled miRNAs, indicating successful transfection (*P*<0.001, Figure 2A). To determine whether cell count reduction was accompanied by morphological changes, we performed PCA of the single-cell profiles. We quantified effects of batch and treatment using ANOVA, finding that batch effects outweighed treatment effects (η² of batch: 0.0505, η² of treatment: 0.0073). Batch effects were corrected using a linear model that first measures the mean deviation per feature and plate among negative control cells and subsequently removes it (Figure 2B, Methods)^30^. This method does not remove all batch effects but reduces the variance they explain by an order of magnitude (η² of batch: 0.0059, η² of treatment: 0.0062). To further mitigate plate-specific effects, we then derived per-plate PCs.

**Figure 2.**
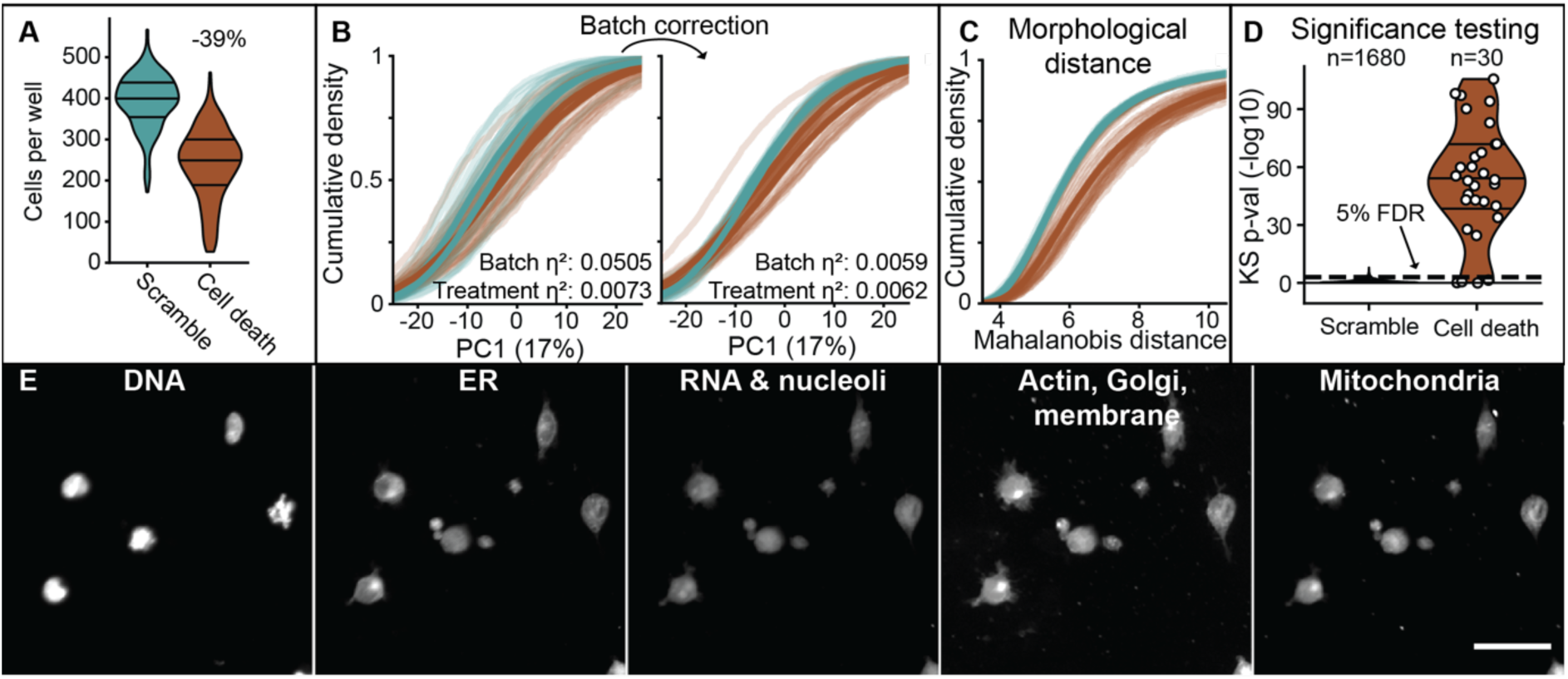
Cell death siRNAs tend to reduce cell counts in HEK293T, compared to cells only treated with scrambled miRNAs, confirming successful transfection. A: Cell counts per well and condition. Horizontal lines on each violin plot indicate 25th, 50th and 75th percentiles. B: Left: At the single-cell profile level, batch effects explain more variance than differences between treatments, as measured by variance explained by each variable. Bold lines represent averages per replicate; thin lines are per-plate profiles. Right: Correcting batch effects with a linear model reduces the differences between replicates and batches, and reduces variance explained by batch by an order of magnitude. C: Single-cell Mahalanobis distances to the control centroid highlight the phenotypic differences of cells treated with cell death siRNAs. D: Left: Background distribution of well-to-well KS tests of negative controls, with the dashed line indicating the 95th percentile. Right: Cell death siRNA treatments induced a significantly different morphology compared to control treatments in 26 of 30 plates tested. E: Per-channel images of HEK293T cells treated with cell death siRNAs. Scale bar is 50 μM.

In this PC space, treated cells were consistently farther from the control centroid (Figure 2C). This showed that the effect of cell death siRNAs on cell morphology was quantifiable and significant at the single-cell level. This effect was reproducible because 26 of the 30 (87%) independently tested plates showed significant differences in cell morphology between the two groups (Figure 2D). Consistent with the computationally inferred change in cell morphological profiles, images of these treated cells show cell rounding and detachment, both features of apoptosis (Figure 2E). In other cell lines, the morphological effects of cell death siRNAs varied, reaching significance in 5%, 10%, 20%, and 68% of experimental replicates for N2a, HepG2, SH-SY5Y, and C2C12 cells, respectively (Supplementary Figure S4) and leading to average cell count reductions of 4%, 27%, 28%, and 17%. This varied response is likely due to the early time point assessed in this study (24 h after transfection), differences in morphological plasticity between cell lines^31–33^, and differences in transfection rate (47%, 26%, 42%, 23%, 53% for HEK293T, N2a, HepG2, SH-SY5Y, and C2C12 cells, respectively, after 48 h).

We also tested an additional positive control that acts independently of transfection, the insecticide rotenone, which impacts tubulin and mitochondria^34,35^. Rotenone yielded substantial morphological changes in HEK293T (87% replicates significant) and HepG2 cells (57%), but less change in other lines (0-14%, Supplementary Figure S5).

Following this confirmation of the sensitivity of our approach to significance testing, we tested its specificity. For this, we investigated miRNAs that are not expected to alter cell morphology. Among all tested miRNAs, 28 have been removed from the most recent version of miRBase (v22.1) following new evidence that they are not encoded in human or mouse genomes, do not map uniquely to the genome, or instead are rRNA or tRNA fragments (Supplementary Table 1). We hypothesised that these retired miRNAs would not alter cell morphology. These 28 retired miRNAs were tested 82 times in our experiments. Using the same hit calling criteria as for other miRNA mimics, only 3 (4%) of these tests were significant at 5% FDR indicating, as expected, that most fail to induce morphological change (Supplementary Table 1).

### Impact of miRNA mimics on cell morphology

We then tested all 2,565 human and 1,900 mouse miRNAs in the library for morphological effects. Of these, 226 (9%) human and 286 (15%) mouse miRNAs were hits in both replicates of one or more cell line (at 5% FDR, Table 1 and Supplementary Table 2). Ten of these human miRNAs induced changes in two human cell lines (Figure 3A). Across organisms, 18 of 4,163 human or mouse sequence-unique miRNAs affected two cell lines, six of these are sequence-identical orthologues, and an additional two were orthologues sharing an identical seed sequence (Figure 3B). Using a different negative control as reference showed agreement with these results (Figure S6). We did not find any miRNAs acting in both replicates of more than two tested cell lines.

**Figure 3.**
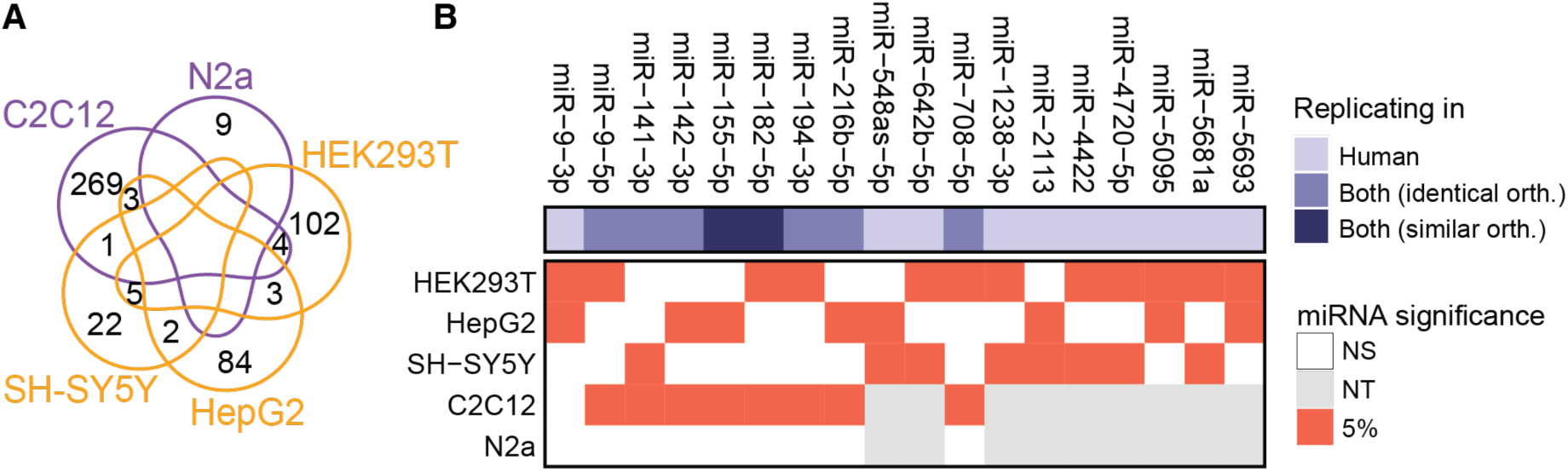
Most miRNA hits are cell line specific, except 18 notable miRNAs. A: Venn diagram showing number of miRNAs being a hit in each cell line, including overlaps. Unlabelled segments indicate no shared miRNAs. B: 18 miRNAs significantly alter cell morphology in both experimental replicates for each of two cell lines, including six orthologues with perfect sequence identity (“identical orth.”) and two orthologues with sequence identity in the seed and up two 2 nucleotide differences (Levenshtein distance) outside the seed (“similar orth.”). NS = not significant, NT = not tested.

**Table 1.** 2,565 human and 1,900 mouse miRNAs, totalling 4,163 miRNAs with a unique sequence, were tested across five cell lines using two experimental replicates. miRNAs that induced significant morphological changes in both replicates were defined as hits. 226 human and 286 mouse miRNAs were a hit in at least one cell line, resulting in 507 unique miRNAs that were hits in at least one cell line.

| Cell line | Hits<br>replicate 1 | Hits<br>replicate 2 | Hits<br>shared | Total sequence-unique miRNA hits by<br>organism | Total unique<br>miRNA hits |
| --- | --- | --- | --- | --- | --- |
| HEK293T | 463 | 262 | 114 | 226 | 507 |
| HepG2 | 353 | 371 | 92 |  |  |
| SH-SY5Y | 262 | 361 | 30 |  |  |
| C2C12 | 787 | 446 | 277 | 286 |  |
| N2a | 112 | 68 | 9 |  |  |

### Morphological effects of miR-155-5p may relate to seed sequence

Among replicating miRNAs was human and mouse miR-155-5p (Figure 3), which has been well characterised in the contexts of inflammation and cancer drug resistance^36–39^. This miRNA significantly altered cell morphology in at least one replicate of four cell lines (Figure 4A). We therefore investigated its impact on each of 142 morphological features relative to miRNAs with seed sequences differing by up to one nucleotide. To do so, we performed t-tests per feature, comparing treated to negative control cells. This resulted in 142 t-statistics per miRNA, representing how much each feature altered following treatment, which we then clustered (Methods). The cellular phenotypic effect conveyed by hsa-miR-155-5p in HepG2 and

**Figure 4.**
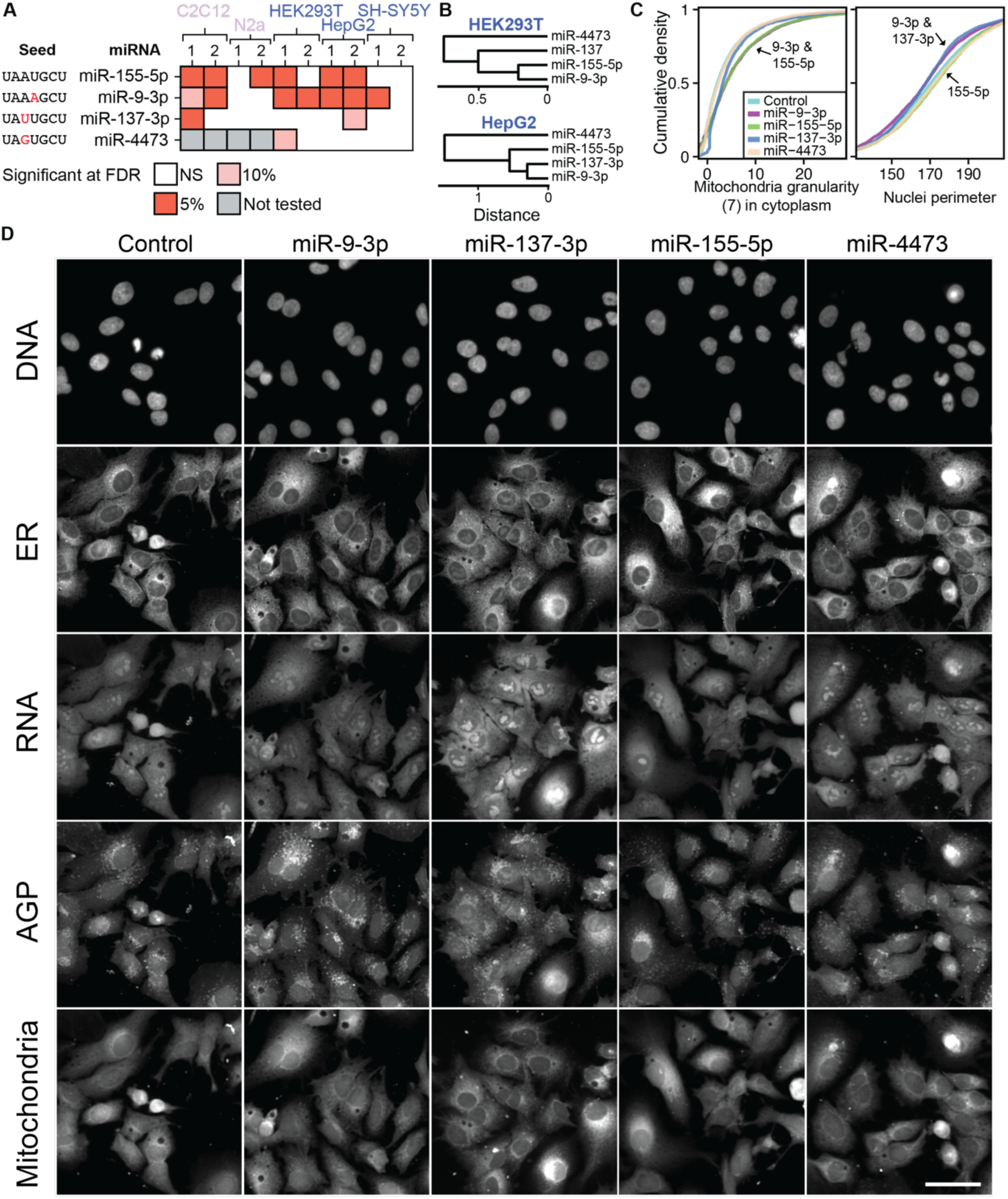
miR-155-5p-induced morphological changes are similar to those induced by seed sequence-similar miRNAs. A: miR-155-5p and three closely related miRNAs affect cell morphology in both human and mouse cell lines. Heatmap shows hits called for replicates 1 and 2 in each cell line. B: Three of the four miRNAs yield similar morphological changes to control cells, resulting in their co-clustering in HEK293T and HepG2 lines. miR-4473, however, does not yield comparable changes. C: In HepG2, miR-9-3p and miR-155-5p impact mitochondrial granularity, and miR-9-3p and miR-137-3p reduce the nuclear perimeter. This view of cell morphological changes helps to explain the clustering observed in B. D: Differences in HepG2 morphology are not immediately discernible by eye, except for differences in vacuole or droplet content. Scale bar is 50 μM.

HEK293T cells was most similar to effects of hsa-miR-9-3p and hsa-miR-137-3p (Figure 4B). Subtle differences in miRNA effects were evident: miR-9-3p and miR-155-5p increased mitochondrial granularity, miR-9-3p and miR-137-3p decreased the nuclear perimeter, and the outgroup, miR-4473, resembled negative control cells of HepG2 (Figure 4C). This fine-grained view helps explain the clustering observed in Figure 4B. The activity of miR-155-5p, miR-9-3p, and miR-137-3p may therefore depend on their seed sequence. Single nucleotide changes in miRNA seed sequence (Figure 4A) are thus associated with similar (hsa-miR-9-3p and hsa-miR-137-3p) and dissimilar (hsa-miR-4473) morphological changes in HepG2 cells. These miRNA-induced changes in morphology were barely discernible by eye, underlining the need for sensitive, quantitative analysis of morphological profiles (Figure 4D).

To connect these cellular changes with molecular phenomena, we next investigated whether these miRNAs – which induce similar cellular changes – share similar mRNA target ontologies. Using gene ontology enrichment (Methods), we found that mRNA targets of miR-9-3p and miR-155-5p were each enriched in TGF-β signaling (both p<0.01), and miR-9-3p’s mRNA targets were associated with mesenchyme development (p<1×10^-5^). MiR-155-5p has previously been implicated in the TGF-β signaling cascade through targeting SMAD proteins, changes to which HepG2 cells are sensitive^36,40–44^. Together, these results show that single-cell analysis can detect subtle changes in cell morphology, similarities in seed sequence can reflect similarities of morphological changes, and cell morphology adds an orthogonal informational resource for miRNA target annotation.

### A shared seed in the hsa-miR-518 family mediates morphological impact

While the above results suggested that a similarity in seed sequence may influence morphological similarity, the case study of hsa-miR-518 shows that seed identity can outweigh miRNA family membership. Of the 11 members of hsa-miR-518, five share an identical seed sequence (Figure 5A). All five miRNAs led to a significant morphological difference (i.e., were all a hit) in HEK293T, whereas the other six miRNAs did not. MiRNAs with the AAAGCGC seed were phenotypically similar and clustered separately from miRNAs with a different seed sequence (Figure 5A). Among the features most changed by these five miRNAs was mitochondrial granularity, which was significantly higher compared to the negative control (Figure 5B). As for hsa-miR-155-5p, this effect is small (Figure 5C). Overall, these findings confirm that identical seed sequences can mediate similar morphological changes.

**Figure 5.**
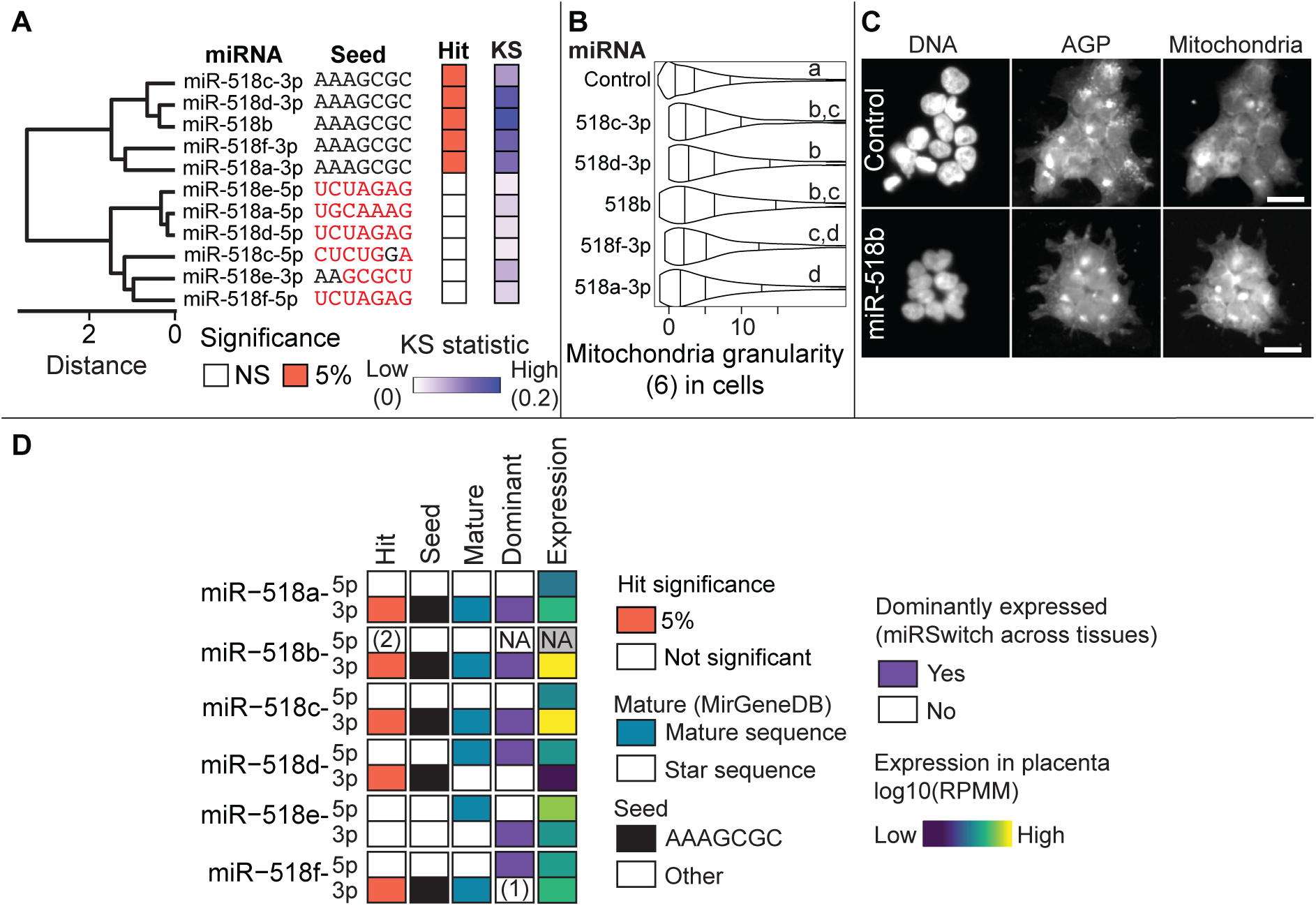
A shared seed in the hsa-miR-518 miRNAs mediates morphological effects in HEK293T. A: The seed AAAGCGC is contained in five of the 11 miRNAs. All five were considered a hit compared to negative control, i.e. significantly altered cell morphology, while the other six did not. Clustering the miRNAs by phenotypic similarity confirms that the five miRNAs induce similar morphological changes (dendrogram). B: Among the top features affected by the five miRNAs is mitochondrial granularity in cells (“Cells_Granularity_6_W5”). Horizontal lines in violins indicate quantiles at 0.25, 0.5 (median), and 0.75. Compared to negative control, the miRNAs have significantly higher mitochondrial granularity. Letters indicate results from pairwise Wilcoxon rank sum tests. Groups that do not share a letter are significantly different at p < 0.05 after Benjamini-Hochberg (BH) correction^51^. C: Images of DNA, AGP of cells treated with negative control or hsa-miR-518b show minor differences in cell morphology. The scale bar is 25 μM. D: miR-518 miRNAs morphological changes are related to seed sequence and endogenous expression patterns. MiR-518a, miR-518c and miR-518f show a consistent pattern of the 3p arm being predominantly expressed in the placenta, the tissue predominantly expressing miR-518^48,50^. MiR-518f switches from 5p to 3p dominance in the placenta specifically, showcasing its tissue specific activity (1). Endogenous expression alone is insufficient to predict morphological activity, however, with miR-518d-3p lowly expressed natively but still showing significant morphological changes as a mimic in our assay, suggesting that the seed sequence is sufficient to induce morphological changes. (2) miR-518-5p is sometimes detected in deep sequencing^47^, but not annotated in miRBase and was not tested in our experiments. Expression in placenta from miRNATissueAtlas^49^.

In vivo, miRNA duplexes are typically processed into the 3p and 5p arms, with the passenger strand (“star sequence”) degraded swiftly while the lead strand (“mature sequence”) can exert its effect of downregulating mRNAs. This imbalance of expressed arms is governed by sequence-specific properties, such as thermodynamics^45,46^. In our experiments, the use of stabilized miRNA mimics allows testing of the passenger strand for biological relevance, too.

Using our hsa-miR-518 case study, we thus tested if morphological impact was related to markers of arm dominance. To this end, we linked our results to information from MirGeneDB^47^, miRSwitch^48^, and miRNATissueAtlas^49^. Endogenous expression of miR-518 members is specific to the placenta^48,50^. Indeed, we find that the arm predominantly expressed in the placenta tends to yield significant morphological changes (Figure 5D). An exception is represented by miR-518d; while the 3p arm contains the shared seed sequence, its 5p arm is annotated as mature and has a higher expression. However, our miRNA experiments show that the seed sequence in its 3p arm still exerts the morphological change. This suggests that expression patterns of miRNA arms are related to but not fully predictive of morphological activity. We stress that here we used miRNA mimics which are chemically stabilized, meaning that hsa-miR-518d-3p is unlikely to exert this effect endogenously. Instead, the research of miRNA mimics using morphology in this way may reveal miRNAs of therapeutic use that would go missed when relying on expression patterns as markers of significance alone.

### Trends of how miRNAs influence cellular morphology

Next, we investigated trends of miRNA seed sequence, annotation confidence, and cell toxicity. Motivated by results from miR-155-5p and miR-518 mimics (Figure 4 and Figure 5), we investigated how morphological changes relate to miRNA seed sequence changes across all miRNAs. Among miRNA hits identified in C2C12, the cell line with the most hits, we found that similar miRNA seeds tended to yield more similar morphological profiles compared to the negative control (Figure 6A, Mann-Whitney-U test, p<0.001 after Benjamini-Hochberg (BH) correction^51^). Identical seeds and seeds differing by one nucleotide induce similar morphologies, but this similarity diminishes as more nucleotide changes are added. This seed sequence-dependent effect mirrors similar findings for shRNAs^24^.

**Figure 6.**
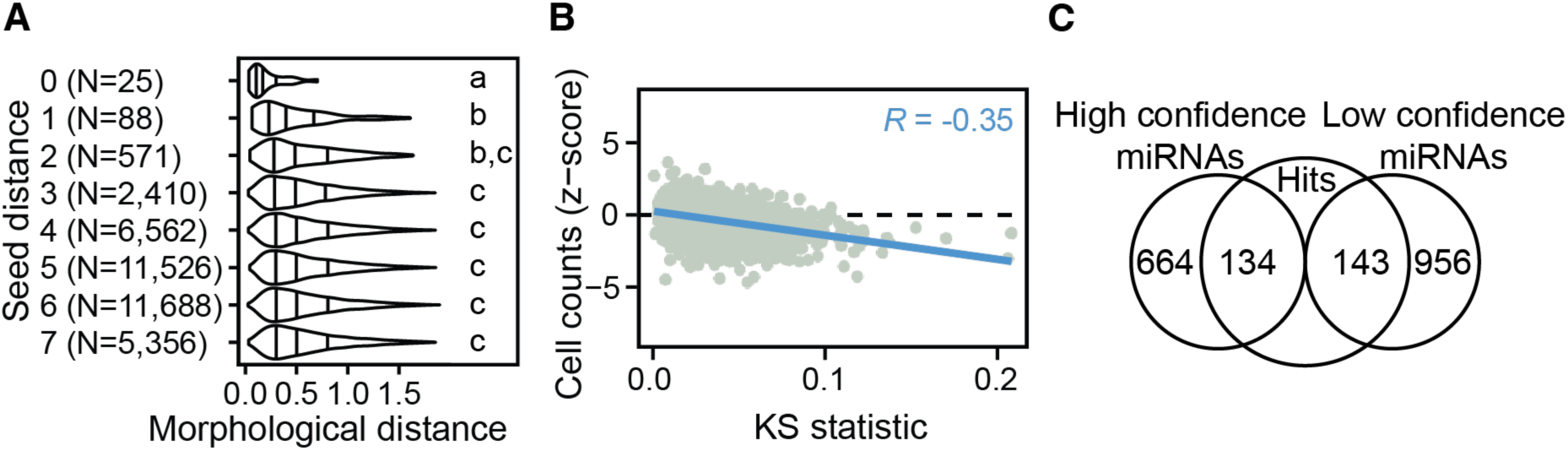
Trends of how miRNAs influence cellular morphology. A: Among C2C12 hits, miRNAs with identical seed sequence yield more similar morphological changes compared to negative controls. Differences between seed sequence were quantified with the Hamming distance. Letters indicate results from Mann-Whitney-U tests between pairs of the 8 groups. Groups labelled with different letters are significantly different at p<0.001 after BH correction. B: Cell count (y-axis) is inversely correlated with cell morphological change (p<0.0001), as measured by the KS statistic (x-axis; Methods). Results are per miRNA and were averaged across all cell lines. C: In C2C12, high confidence miRNAs were more frequently significantly different (134/801) from controls than non-high confidence miRNAs (143/1099), a difference which is significant (Chi-square test p<0.05).

Differences in morphological features are negatively correlated with cell counts, indicating that cytotoxicity is a predictor of morphological change^52^. In our miRNA experiments, we also observed a negative correlation between cell counts and morphological difference to negative control, although this effect varied by cell line (Figure 6B and Supplementary Figure S7). However, one may find miRNAs such as miR-19a-5p escaping this trend by mediating large morphological changes without altering cell count (Supplementary Figure S7). As observed in our positive control experiments using cell death siRNAs, cytotoxicity is therefore associated with morphological changes among surviving cells.

We finally investigated whether well-studied miRNAs tended to mediate stronger morphological changes. miRBase annotates a subset of miRNAs as “high confidence” based on experimental evidence, including their processing by Drosha and Dicer proteins^6^. Of miRNAs tested in this experiment, 861 (34%) human and 801 (42%) mouse miRNAs are annotated as high confidence. In C2C12, 17% (134 of 801) of “high confidence” miRNAs were a hit in both replicates (Figure 6C), significantly higher than this proportion among other miRNAs (13%; 143 of 1099; Chi-square test, p < 0.05). This indicates that mimics of miRNAs with experimental evidence are more likely to yield a discernible cellular morphological response.

## Discussion

We comprehensively profiled the morphological effects of 2,565 human and 1,900 mouse individual miRNA mimics on single cells for three human and two mouse cell lines. Of these, 226 human and 286 mouse miRNAs had a significant impact on cell morphology (Table 1), including 18 miRNAs in multiple cell lines (Figure 3). We envision that this comprehensive reference of single-cell morphological profiles upon miRNA treatment of diverse cell lines will begin to bridge the current gulf in understanding between miRNA cellular and molecular function.

Seed sequence-similar miRNAs can mediate similar changes in cell morphology (Figure 4, Figure 5, Figure 6A). miR-155-5p is expected to induce morphological effects similar to miRNAs that differ slightly in the seed sequence. This is because about 40% of miR-155-5p targets are bound with a mismatch or wobble in the seed sequence, most commonly in seed positions 4 and 6^53^. Nevertheless, it was unexpected that miR-155-5p was more similar to miR-137-3p than miR-9-3p in affecting HepG2 cell morphology, because miR-137-3p differs in sequence elsewhere, at seed position 3. For hsa-miR-581 members, we observed a strong similarity among miRNAs sharing the same seed, confirming the paradigm of the seed sequence’s crucial role.

Our understanding of miRNAs remains limited due to: i) cell type specificity of their expression^21^, ii) cell type specific expression of their target mRNAs, iii) imperfect miRNA annotation^17,18,54^ and, iv) inconsistencies in the miRNA literature^2,55^. While much research has sought to quantify miRNA abundance in various tissues, fewer studies have systematically linked miRNAs to phenotypes. Also, those studies that were systematic often focused on specific outcomes, such as cell proliferation, rather than measuring diverse phenotypes^56–58^. Our large-scale single-cell study of miRNA effects on cellular morphological phenotypes begins to overcome these limitations.

Single-cell analysis accounts for: i) incomplete transfection, because not all cells are transfected and thus are expected to display altered phenotypes, ii) variability within cell lines, because averaging profiles obscures dynamic processes such as mitosis^46^, and iii) the possibility of miRNA-induced subpopulations, for example if miRNAs were to induce cell cycle arrest. Our single-cell framework for hit calling identifies miRNAs that cause significant morphological changes, in contrast to previous single-cell research which predicted mechanism-of-action and dose-dependent phenotypic changes^46^. Further, the framework reduces batch and plate-position effects through rigorous batch correction and feature selection, thereby minimising technical artefacts. In these applications, our analysis methods retain known biological differences, while allowing discovery of morphological changes for subsets of cells. We anticipate that these single-cell analysis methods, consolidated in the Python software scmorph^29^, prove useful for other complex morphological profiling experiments when subpopulation-level changes are likely. Among the most obvious applications are experiments with multiple cell lines^59,60^, differentiating cells^61^, and transfected treatments (as in this work). In these applications, our analysis methods retain known biological differences, while allowing discovery of morphological changes of a subset of cells.

Mullokandov et al. found that only the most highly expressed miRNAs suppressed their target genes^62^. In contrast, our findings show that, even at lower concentrations, hundreds of miRNAs induce morphological change. This may be attributable to miRNAs conveying small but coordinated effects on many mRNA targets, or to them regulating the abundance of transcription factors, thereby changing the transcription landscape more widely^14,63^. Further, we showed that cytotoxic miRNAs induce stronger morphological shifts, mirroring similar results in small molecule screens (Figure 6B)^60^. This suggests that miRNAs may mediate morphological and cytotoxic effects at relatively modest concentrations.

Overall, across diverse cell lines, most miRNAs do not yield significant morphological changes. This is consistent with results that over 60% of miRNAs do not discernibly suppress their mRNA targets, as measured with a reporter gene linked to miRNA target sites^62^. Compared to simply counting cells, we identified miRNAs that alter cell morphology without causing cytotoxicity (Supplementary Figure S7). Our approach provides several advantages over previous large-scale studies of miRNA activity that focused on single cell lines, such as HeLa^56^, or on a specific outcome, most often cell proliferation or viability^56–58^, even when using high-content microscopy^64,65^. Instead, we identified significant morphological changes across five cell lines, including miRNAs that did not impact cell viability but still changed cell morphology. We speculate that hit miRNAs may serve as interesting leads for further research, as they yield cellular phenotypes beyond changes in gene expression.

In line with previous findings by others^33,66^, we found that cell lines responded heterogeneously to perturbations: HEK293T readily changed morphology after perturbation with positive controls and any of over 100 different miRNAs. Other cell lines had more variable responses. Differences in transfection efficiency and cell cycle time may explain some of the observed differences. Equally, endogenous miRNA expression may influence the observed activity of transfected miRNAs – a transfected miRNA may not induce a shift in phenotype if it is already highly expressed natively.

Our approach is limited by only considering miRNAs to be significant by their induction of measured morphological changes, disregarding other cellular effects and molecular changes such as mRNA levels which could provide complementary information^67,68^. Ideally, RNA and morphology of the same cell could be quantified, yet this remains experimentally challenging. A complement to this study would be a morphological profiling experiment of miRNA knockouts, which is achievable due to recent advances^69,70^. An additional limitation is that during hit calling we prioritised true positives over false negatives through stringent significance testing, leading us to underestimate the true number of cell morphology-associated miRNAs. Lastly, we performed the Cell Painting assay after 24h to avoid overly dense growth of cells in the microwells, which could have masked morphological changes. Later time points (e.g. 48-72h) would likely lead to stronger effects^31,32^. The number of miRNAs with a significant effect on morphology at later time points would therefore be expected to be higher than found here.

This study provides a resource to help assess whether a particular miRNA affects cell morphology in any one of 5 cell lines (Supplementary Table 2) and which features it affects most (Supplementary Table 3). For more systematic miRNA research, our whole single-cell dataset can be intersected with external information on miRNAs, such as their effect on the transcriptome, or with data on miRNA-binding dynamics of target mRNAs^71^. In summary, we anticipate that this study will serve as a starting point for future systematic characterizations of miRNAs.

## Methods

### Reagents and tools table

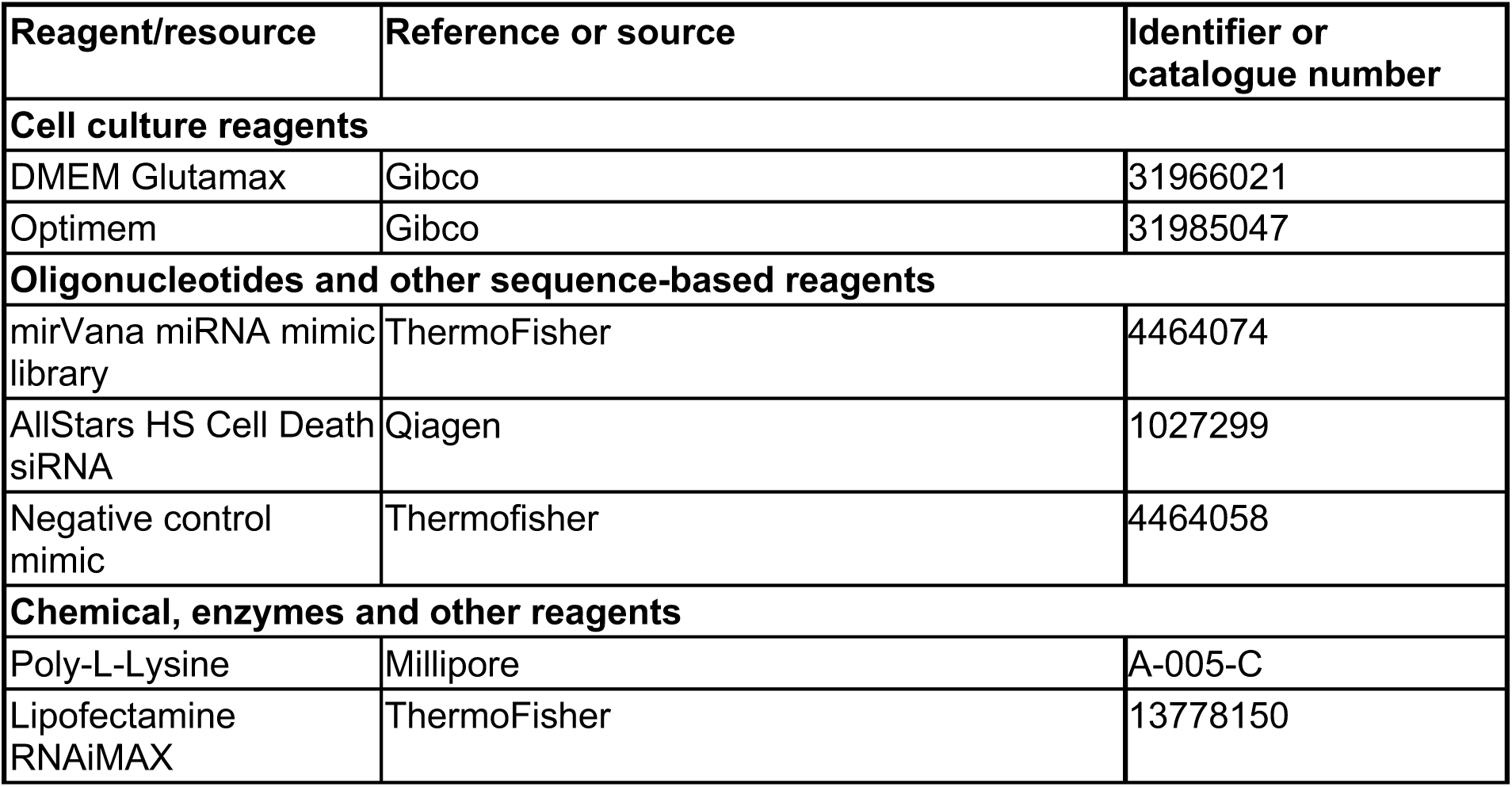
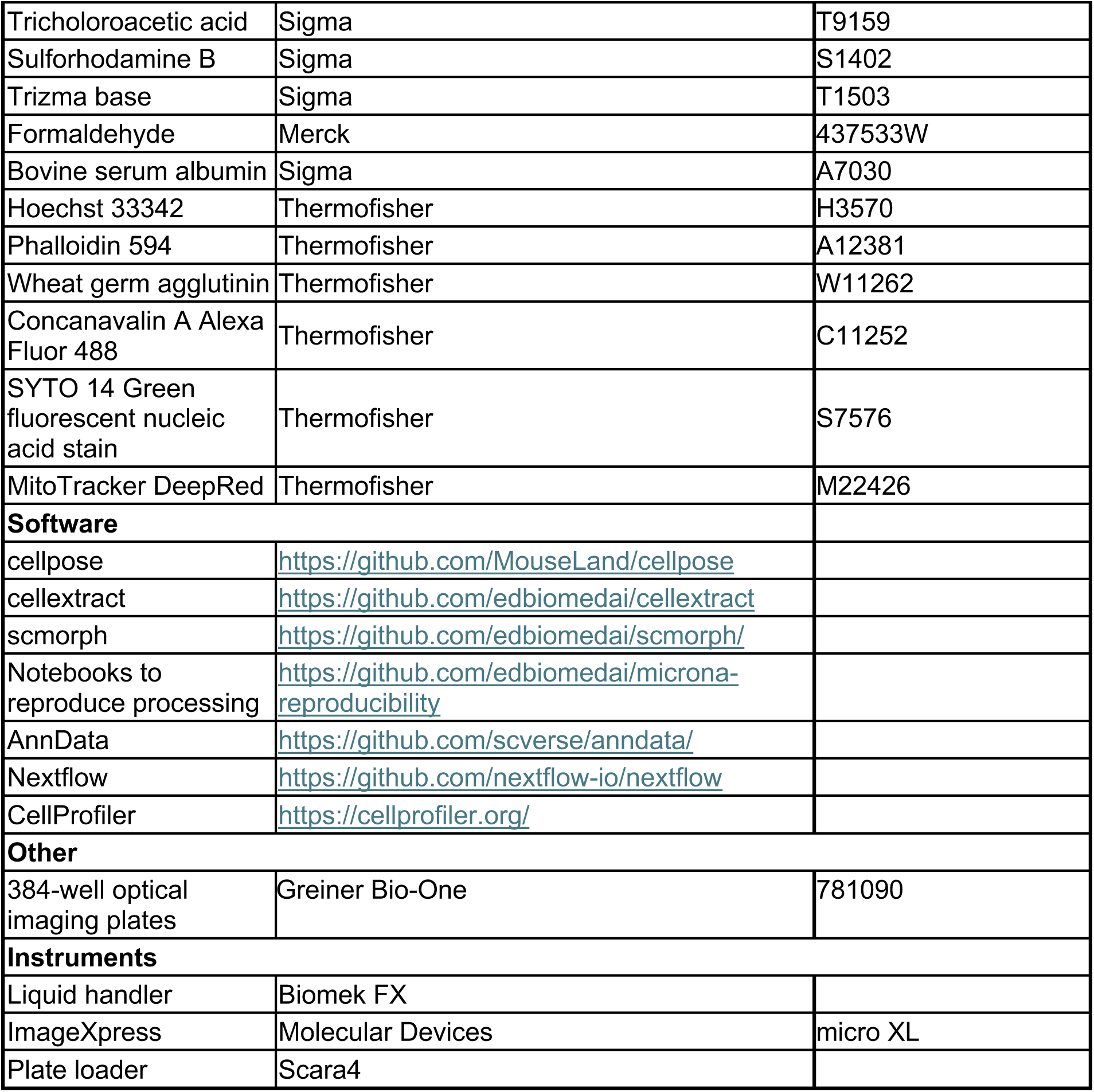

| Reagent/resource | Reference or source | Identifier or catalogue number |
| --- | --- | --- |
| **Cell culture reagents** |  |  |
| DMEM Glutamax | Gibco | 31966021 |
| Optimem | Gibco | 31985047 |
| **Oligonucleotides and other sequence-based reagents** |  |  |
| mirVana miRNA mimic library | ThermoFisher | 4464074 |
| AllStars HS Cell Death siRNA | Qiagen | 1027299 |
| Negative control mimic | Thermofisher | 4464058 |
| **Chemical, enzymes and other reagents** |  |  |
| Poly-L-Lysine | Millipore | A-005-C |
| Lipofectamine RNAiMAX | ThermoFisher | 13778150 |
| Trichloroacetic acid | Sigma | T9159 |
| Sulforhodamine B | Sigma | S1402 |
| Trizma base | Sigma | T1503 |
| Formaldehyde | Merck | 437533W |
| Bovine serum albumin | Sigma | A7030 |
| Hoechst 33342 | Thermofisher | H3570 |
| Phalloidin 594 | Thermofisher | A12381 |
| Wheat germ agglutinin | Thermofisher | W11262 |
| Concanavalin A Alexa Fluor 488 | Thermofisher | C11252 |
| SYTO 14 Green fluorescent nucleic acid stain | Thermofisher | S7576 |
| MitoTracker DeepRed | Thermofisher | M22426 |
| **Software** |  |  |
| cellpose | https://github.com/MouseLand/cellpose |  |
| cellextract | https://github.com/edbiomedai/cellextract |  |
| scmorph | https://github.com/edbiomedai/scmorph/ |  |
| Notebooks to reproduce processing | https://github.com/edbiomedai/microna-reproducibility |  |
| AnnData | https://github.com/scverse/anndata/ |  |
| Nextflow | https://github.com/nextflow-io/nextflow |  |
| CellProfiler | https://cellprofiler.org/ |  |
| **Other** |  |  |
| 384-well optical imaging plates | Greiner Bio-One | 781090 |
| **Instruments** |  |  |
| Liquid handler | Biomek FX |  |
| ImageXpress | Molecular Devices | micro XL |
| Plate loader | Scara4 |  |

### miRNA mimic library

A mirVana miRNA mimic library containing all human and all mouse miRNAs (miRBase v21) was obtained (ThermoFisher, 4464074, 0.25 nmol). This encompassed 2,565 human and 1,900 mouse miRNA mimics (called “miRNAs” throughout for brevity). Handling of the miRNA library used an automated liquid handler (Biomek FX). The library was prepared first as a 5 μM stock in DNase/RNase free dH_2_0, before creating intermediate plates with 2 μL miRNA stock in 59 μL dH_2_0. miRNAs for transfections were taken from intermediate plates. miRNA sequences were obtained from miRBase (v21) and seed sequences were defined as positions 2-8.

### Cell culture

All cell lines were grown in DMEM Glutamax (Gibco, 31966021) supplemented with 10% fetal calf serum, incubated at 37°C with 5% CO_2_. Cell numbers were optimised for 384-well optical bottomed imaging plates (Greiner Bio-One, 781090). Prior to cell seeding, imaging plates were coated with Poly-L-Lysine (Millipore, A-005-C) to ensure adherence of cells. Numbers of cells seeded per well were: 1600 cells/well for HEK293T, 1500 cells/well for SHSY-5Y, 1900 cells/well for HepG2, 1000 cells/well for N2a and 900 cells/well for C2C12.

### Cell transfection

Cells were transfected with the miRNA mimics via reverse transfection. For each cell line a transfection mix was made from an optimised concentration of Lipofectamine RNAiMAX (ThermoFisher, 13778150) in Optimem (Gibco, 31985047), using 12 μL RNAiMAX per 1 ml Optimem for HEK293T, 18 μL for SH-SY5Y and C2C12, and 24 μL for HepG2 and N2a. 5 μL of this transfection mix was added to each well of the 384-well imaging plates. 2 μL of the miRNA mimic stock was then added into the transfection mix and the plates vortexed. The plates were left to incubate at room temperature for 30 min before addition of 25 μL of cells in culture medium for a final miRNA mimic concentration of 10 nM. Note that the final intracellular concentration of functional miRNAs depends on transfection efficiency and processing of vesicles and will therefore vary across cell lines and conditions^72^. Plates were transfected in batches of up to 4 plates to ensure the timing of cell addition was optimal for transfection efficiency. Cells were incubated for 24 hours prior to fixing.

Transfection rates were assessed using the SRB assay for cytotoxicity^73^. For this, HEK293T was incubated with 0.06 μL/mL RNAiMAX, whereas concentrations for other cell lines were 0.09 μL/mL for SH-SY5Y and C2C12, and 0.12 μL/mL for HepG2 and N2a. Cell death siRNAs were added and incubated for 48 h. Then, cells were fixed using tricholoroacetic acid (T9159, Sigma) and stained with SRB (S1402, Sigma). Excess dye was washed away with acetic acid after 30 min, and cell-bound dye was redissolved using trizma base. Optical density was measured at secondary peak of 490 nm, with results from siRNA-treated cells compared to cells in cell medium.

### Experimental design

The miRNA library was profiled in three human (HEK293T, SH-SY5Y and HepG2) and two mouse cell lines (C2C12 and N2a). Within each cell line, each miRNA was profiled in two wells (same plate, separate well) and two replicates (separate experimental batch and plate). During analyses, the two wells on the same plate were pooled. miRNAs were split over 15 plates for human and 11 plates for mouse cell lines (cf. Figure 1B for plate layout). Each plate included multiple wells each of three negative and two positive controls: DMSO only, DMSO with transfection mix, and DMSO with negative control mimic (i.e. scrambled miRNA, Thermofisher, 4464058) for negative controls, and AllStars HS Cell Death siRNA (Qiagen, 1027299) and 5 nM rotenone for positive controls. The positive controls were chosen to investigate transfection efficiency and changes to mitochondrial networks, respectively. Throughout the study, the main negative control referenced was scrambled mimics because it controls for the effect of miRNA mimics being loaded into RISC and the potential competition with endogenous miRNAs. We provide results from the contrast of miRNA mimics and DMSO with transfection mix in Supplementary Tables 4-6 for completeness.

### Cell staining and imaging

We followed the Cell Painting protocol as described by Bray et al. (2016)^23^, with changes only to the mitochondrial staining procedure. Briefly, cells were fixed 24 h after transfection of miRNA mimics by addition of an equal volume of 9% formaldehyde (Merck, 437533W) to give a final concentration of 4% formaldehyde. Cells were incubated in this solution for 30 minutes at room temperature and then washed twice with PBS. Cells were then permeabilised for 20 minutes at room temperature with 50 μL per well of 0.1% Triton-X in a solution of 1% bovine serum albumin (BSA) (Sigma, A7030) in PBS. A Cell Painting solution was made up with dyes as listed in Table 2 in 1% BSA/PBS. Of this solution, 20 μL was added to each well and incubated away from light for 30 minutes. Wells were then washed three times with PBS and the plates sealed and stored in the dark at 4°C until imaging.

**Table 2.** Fluorescent stains used for Cell Painting, cellular targets and imaging channels. Wavelength ex/em = excitation and emission, respectively.

| Stain | Target | Wavelength<br>(ex/em [nm]) | Channel | Assay<br>concentration<br>[μL/mL] | Cat. No. |
| --- | --- | --- | --- | --- | --- |
| Hoechst 33342 | Nuclei | 387/447 | DAPI | 0.2 | H3570 |
| Phalloidin 594 | F-actin | 562/624 | TxRED | 0.14 | A12381 |
| Wheat germ<br>agglutinin | Golgi and<br>plasma<br>membrane | 562/624 | TxRED | 1 | W11262 |
| Concanavalin A<br>Alexa Fluor 488 | Endoplasmic<br>reticulum | 462/520 | FITC | 4 | C11252 |
| SYTO 14 Green<br>fluorescent nucleic<br>acid stain | RNA, nucleoli | 531/593 | SYTO14 | 0.6 | S7576 |
| MitoTracker<br>DeepRed | Mitochondria | 628/692 | CY5 | 0.6 | M22426 |

Imaging was carried out on an ImageXpress micro XL (Molecular Devices, USA) equipped with a robotic plate loader (Scara4, PAA, UK). Four images were captured per well with 20x magnification and 5 fluorescent channels (see Table 2). Exposure times were optimised separately for each cell line but kept consistent across the plates and replicates for each cell line.

### Cell segmentation

To detect cells from the 205,824 images we developed an approach based on CellPose^74^, a deep-learning model for cell and nuclei segmentation, that accounts for diverse cell shapes, multinucleation and segmentation errors. Briefly, it first detects cell boundaries from the TxRed channel and in a second pass detects nuclei within these cells. More specifically, cell boundaries were detected from images with the cyto2 CellPose model, with expected sizes listed in Table 3. Cells touching image boundaries were removed from further analysis. To detect nuclei within the cell boundaries, single-cell crops were created with pixels not belonging to the targeted cell set to the background pixel value 0. Crops were padded with a 20 px boundary on each side to ensure the presence of sufficient background for CellPose auto-normalization. Another round of segmentation of the DAPI channel to detect nuclei using the cyto CellPose model was then conducted for each cell crop. The resulting nuclei predictions were trimmed to ensure that detected nuclei fell strictly within the detected cell boundary. Note that this method may detect more than one nucleus per cell, as is the case for multinucleated, mis-segmented and mitotic cells. In this study, only mono-nucleated cells were analysed, because multinucleated cells tended to be either mis-segmented or M-phase cells prior to cytoplasmic division. Nuclei masks were then gathered across crops to match the original image dimensions. Lastly, cytoplasmic masks were created that represent cell masks with the nuclear portion removed. These and other analytical steps until and including feature extraction were facilitated by a Nextflow pipeline found at https://github.com/edbiomedai/cellextract. Together, these steps yield a cell, nuclear, and cytoplasm mask per image.

**Table 3.** CellPose size parameters during cell and nuclei segmentation.

| Expected size (μM) | HEK293T | HepG2 | SH-SY5Y | C2C12 | N2a |
| --- | --- | --- | --- | --- | --- |
| Cells | 17 | 30 | 17 | 34 | 17 |
| Nuclei | 10 | 15 | 10 | 17 | 10 |

### Image correction and QC

Fluorescent microscopy images frequently contend with uneven illumination within and across images^25^. While the employed microscope integrated field-of-view illumination correction, we noticed that, across images, background intensities varied, possibly due to minor differences in focus. We aimed to remove the average background intensity from all pixels to ensure equal measurements across all images. To this end, per image and channel, the 25^th^ percentile of background pixels was subtracted from all pixels in the image. We chose the 25^th^ percentile because background detection relies on good cell segmentation. Choosing a higher percentile might therefore risk the inclusion of incorrectly segmented foreground when estimating background intensities. After this correction step, pixels with negative intensities were set to zero intensity, indicating background. Next, 78 image QC features were computed using CellProfiler’s MeasureImageQuality module, a description of which can be found in the CellProfiler manual^75^ and in Bray et al. (2012)^76^. Features included object counts, descriptors of image-level intensities and measures of focus. These well-established metrics can capture a variety of quality issues, including precipitate and air bubbles.

Image QC was then performed using a kNN-based metric. The underlying assumption of this approach was that images of low quality will result in outlier QC features (measured above) and that detection of such outliers can therefore remove these images. Specifically, image QC was performed per replicate by centre-scaling image QC features and computing the first three principal components (PCs) from the 78 QC features. For each data point, i.e. image, in this PC space, we then measured the distance to the 10^th^ nearest neighbour, arguing that images of lower quality will have a lower neighbourhood density and thus larger distance to neighbouring images. The image QC distance was defined as 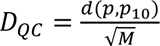, where *d*(*p*, *p*_*+_) is the Euclidean distance of an image *p* to its 10^th^ nearest neighbour, *p*_*+_ and M is the number of PCs the distance *d* is measured in. Images with a distance greater than 0.1 (arbitrary units) were flagged and removed from further analysis. This QC step removed ∼1.1M cells (∼5%, Supplementary Figure S1). Without performing the image QC step, the number of significant hit miRNAs changes from 226 (with QC) to 334 (no QC) in human and from 286 (QC) to 263 (no QC) in mouse cell lines. Image QC, like other processing steps taken, can be reproduced using the set of reproducibility notebooks found at https://github.com/edbiomedai/microna-reproducibility.

### Single-cell feature extraction

Single-cell features were computed from images with matched nuclei, cell and cytoplasm mask using the scikit-learn and skimage libraries, with granularity features computed via CellProfiler. Features describing cell size and shape, channel intensities, distribution, texture and granularity were computed for 1,386 features per cell. Features missing valid measurements in >90% of cells were removed from analysis, leaving 1,377 features (Supplementary Figure S1, Supplementary Table 7). We thus derived features for 22,999,877 cells, exporting results in AnnData format^77^.

### Batch correction

To assess the magnitude of batch and treatment effects of cell death siRNAs in HEK293T (as in Figure S2B), we tested the proportion of variance in position on PC1 explained by treatment and batch (plate number) using ANOVA. The η² statistic was used to measure the variance explained by treatment and batch.

We used a linear model, which maintains interpretability of features, to reduce inter-plate variability (hereafter termed batch effects). Following Cole et al.^30^, who developed a linear model that can remove nested batch effects (equation 3 in Cole et al.^30^), we computed the average batch effect per feature *i* as

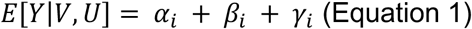

where *V* and *β* are the model matrix and coefficient of known factors of wanted variation, e.g. cell line differences, *U* and γ are the model matrix and coefficient of sources of unwanted variation such as batch effects, and α is an intercept. Since the model is built across all single cells *n*, the matrices *Y*, *V* and *U* all have *n* rows and respectively *J*, *M*, and *H* columns, containing one-hot encoded information of the covariates. This model is fit to negative control cells across batches, which allows identification of the batch effect γ. Subsequently, batch correction is performed for all single-cell profiles, treated or not, by removing the average batch effect *X*^<^, = *X*, − *γ*,. Note that the nested model can maintain differences between cell lines in the form of the *β* coefficient. However, we did not make use of this option in the present study, because we did not directly compare cell morphologies between cell lines. Instead, we performed batch correction for each cell line and replicate experiment separately (Supplementary Figure S1). This method and others used for downstream analysis were implemented in scmorph, our single-cell morphological profiling package in Python^29^.

### Removing plate map effects

After batch correction, we observed considerable association of features with well positions within a plate. We tested each feature for association with column or row position as well as plate ID using a Kruskal-Wallis H-test per replicate and cell line. Note that this test was performed on all wells including treated wells, because negative controls were not randomly dispersed within plates. This means that features may incorrectly be associated with technical variability when strong biological effects are associated with e.g. column position. This is mitigated by testing across all plates in a replicate experiment, making individual treatments less likely to have undue influence on the statistic. For each covariate *j* (column, row, or plate), we compute the *H* statistic for each feature *X*,, yielding a test statistic *H*_,-_ for each such combination. The collection of all test statistics *H*_,-_ across all features *i* is referred to as *H*^?⃑^_-_. Then, we retain only features that satisfy

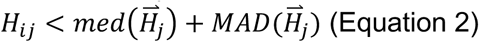

in each cell line and replicate, where *med* is the median and *MAD* is the median absolute deviation. Across all cell lines and replicates, 142 features never exceeded this threshold and were therefore retained for further analysis (Supplementary Table 7, Supplementary Figure S1). Features related to MitoTracker and SYTO 14 Green were more often removed than features from other dyes, highlighting potential issues with experimental procedures (Figure 7). This is in line with previous observations that measures of fluorescence, including SYTO 14, are more often affected by positional effects^78^. Only these 142 features not associated with confounders were retained in further analyses.

**Figure 7.**
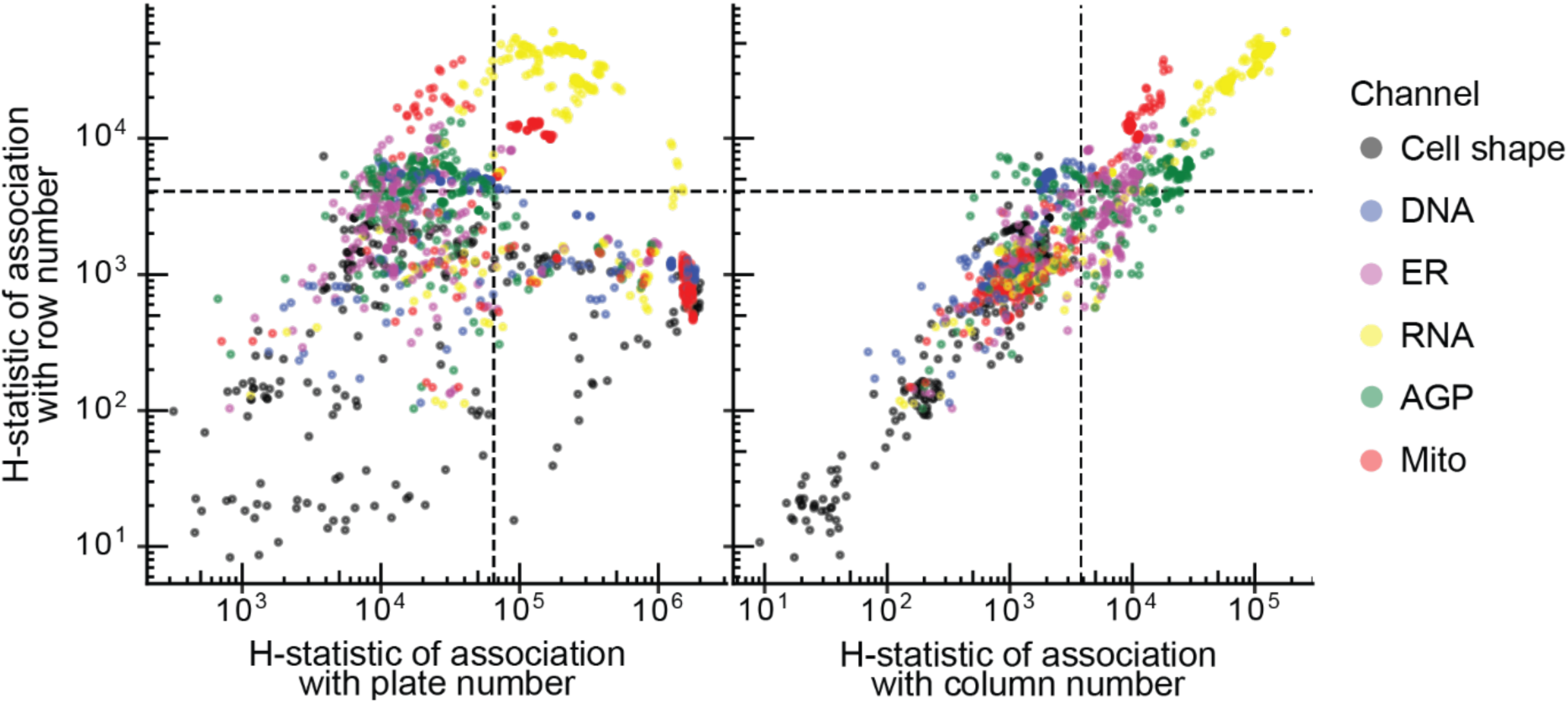
For each feature, represented by a point, the Kruskal-Wallis (KW) test of association with known covariates in HEK293T replicate experiment 1, including plate positions (row, column, left panel) and batch effects (PlateID, right panel y-axis) was computed. MitoTracker and SYTO 14 were more often associated with technical effects. Features measuring cell shape and size were the least frequently discarded. Dashed lines indicate the cutoff for each covariate above which features were discarded.

### Hit calling

To identify treatments that change morphology compared to a negative control, a PCA-based hit calling pipeline was devised. Hit calling was performed strictly per plate to reduce any residual impact of inter-plate differences. First, for each plate, the cell × feature matrix of morphological profiles was standardized (z-scored) and used to compute a PCA, retaining the first 10 PCs (Supplementary Figure S8). Second, the centroid of negative control cells in each PC was obtained. Third, the control cells’ covariance matrix in PCA space was estimated. Fourth, using the inverse of this covariance matrix, the Mahalanobis distance from each cell to the centroid was computed as:

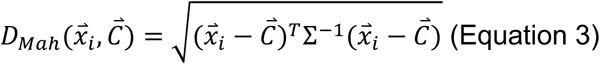

Here 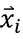, and 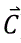 are the PCA-coordinate vector of the *i*th cell and negative control centroid, respectively, and Σ^1^* is the inverse covariance matrix estimated from the PC coordinates of negative control cells. Fifth, for each treatment including negative control, all single-cell distances to the centroid were used to build an empirical cumulative distribution. Sixth, the Kolmogorov–Smirnov (KS) statistic between ECDF from a treatment and the negative control ECDF was computed. To assess statistical significance of observed morphological changes, a null distribution was created by repeating steps two to six, retaining only negative control cells and defining wells as treatments. By using each negative control well as a reference well and computing the well-to-well KS statistic, an estimate of the heterogeneity present within negative control cells was created. False discovery rate (FDR) was controlled per plate by defining the 5^th^ percentile of p-values of KS tests from the null distribution as significance threshold. Treatments were considered hits if the p-value of the KS test comparing the ECDF of treated cells to negative control cells was lower than this threshold in both experimental replicates. This pipeline was applied to each plate independently to further reduce the impact of residual batch effects, with results being aggregated on the level of KS statistics after controlling for false-discovery rate using the Benjamini-Hochberg method^51^.

### Sensitivity of hit calling depends on population size

We use the KS statistic and its p-value to assess significance of morphological changes. As described above, we create a background distribution by performing pairwise tests between negative control wells. The population size in these tests is the number of cells in each well, limiting the lower bound of p-values achievable in the background distribution (Figure 8). When testing treatments, wells on the same plate are pooled, thus increasing the population size and increasing power. This means that a treatment with a KS statistic of e.g. 0.2 may be considered significant even if a KS statistic of 0.25 was considered not significant in the background distribution. However, the lower bound of the p-value asymptotically approaches zero with sample sizes below the number of cells typically observed per well in negative controls (320-810 for SH-SY5Y and HepG2, respectively) (Figure 8). Given these sample sizes, it is unlikely that the lower bound of the p-values is restricting our power to detect differences between control wells, and we can therefore assume that our null distribution accurately captures morphological heterogeneity naturally present in these experiments. That said, at an expected KS statistic of 0.2, sample size impacts p-values by orders of magnitude (Figure 9). It should therefore be assumed that the realised FDR is higher than the intended α. To mitigate this problem, we additionally require treatments to be considered significant in both replicates to be termed a hit.

**Figure 8.**
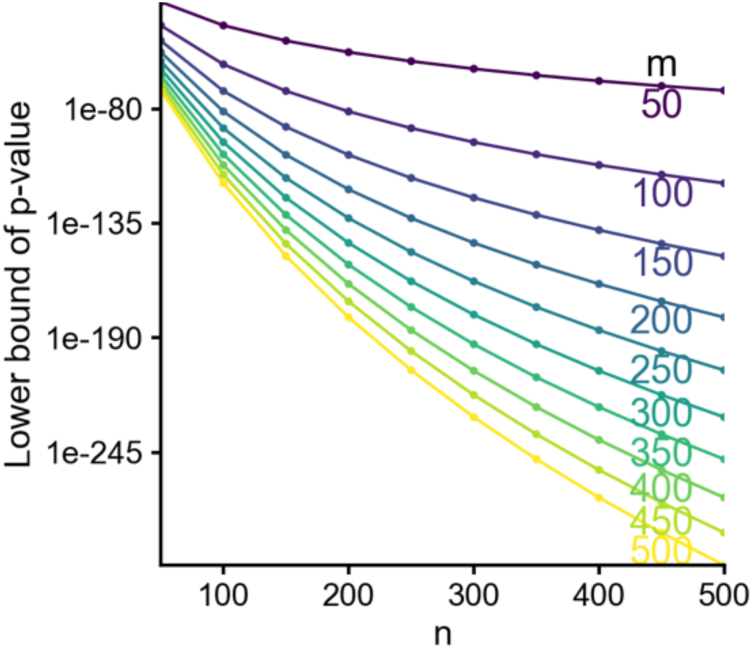
The lower bound of the p-value of a two-sample two-sided KS test is determined by the sample sizes of both samples, n and m.

**Figure 9.**
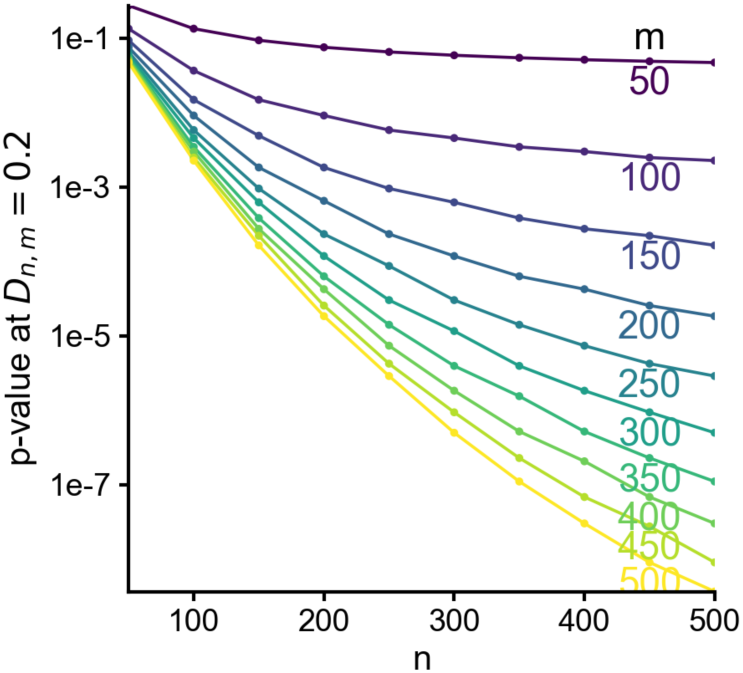
Sensitivity analysis of the KS test p-values, demonstrating that sample sizes of the two samples (n and m) impact expected p-value by orders of magnitude.

### Morphological profile clustering

The outlined hit calling methodology operates on PCs and is therefore not readily interpretable. An orthogonal analysis therefore compared morphological profiles by computing the per-treatment and per-feature t-statistic between treated and negative control cells. For each treatment, cell line and replicate, a vector of 142 t-statistics describing per-feature differences was thus derived. When comparing treatments, as in Figure 4B, the Spearman correlation coefficient between two treatments’ t-statistic vectors was used as a descriptor of similarity. Hierarchical clustering with average linkage and Euclidean distance was then used to cluster treatments.

### miRNA target enrichment

Predicted miRNA targets were obtained from TarBase^79^ and filtered for interaction scores greater than 0.7. Only expressed mRNA targets were considered, which were defined as genes with at least 1 FPKM expression in baseline expression data. Baseline gene expression data for HepG2 was obtained from Gene Expression Omnibus with accession number GSM3478960. Ontology enrichment of targets was tested using g:Profiler^80^, using as background all expressed genes (>=1 FPKM) and with KEGG and all GO terms as sources.

## Data availability

All microscopy images have been uploaded to the Cell Painting Gallery under the prefix cpg0046-microrna^81^. Various pre-processed data, including the cell × feature matrix used in all analyses, are also available through the Cell Painting Gallery.

## Funding and acknowledgments

We thank John C. Dawson for assistance in experimental procedures and Neil O. Carragher for providing reagents and equipment. We also thank Narongrit Sutiyaporn and James Kelly for testing the code and verifying reproducibility of our hit calling results. JW is supported by an MRC PhD studentship (grant no. MC_ST_00035). This work was funded by the Wellcome Trust (grant no. 106956/Z/15/Z). For the purpose of open access, the author has applied a Creative Commons Attribution (CC BY) licence to any Author Accepted Manuscript version arising from this submission.

## Supplementary Figures

**Figure S1.**
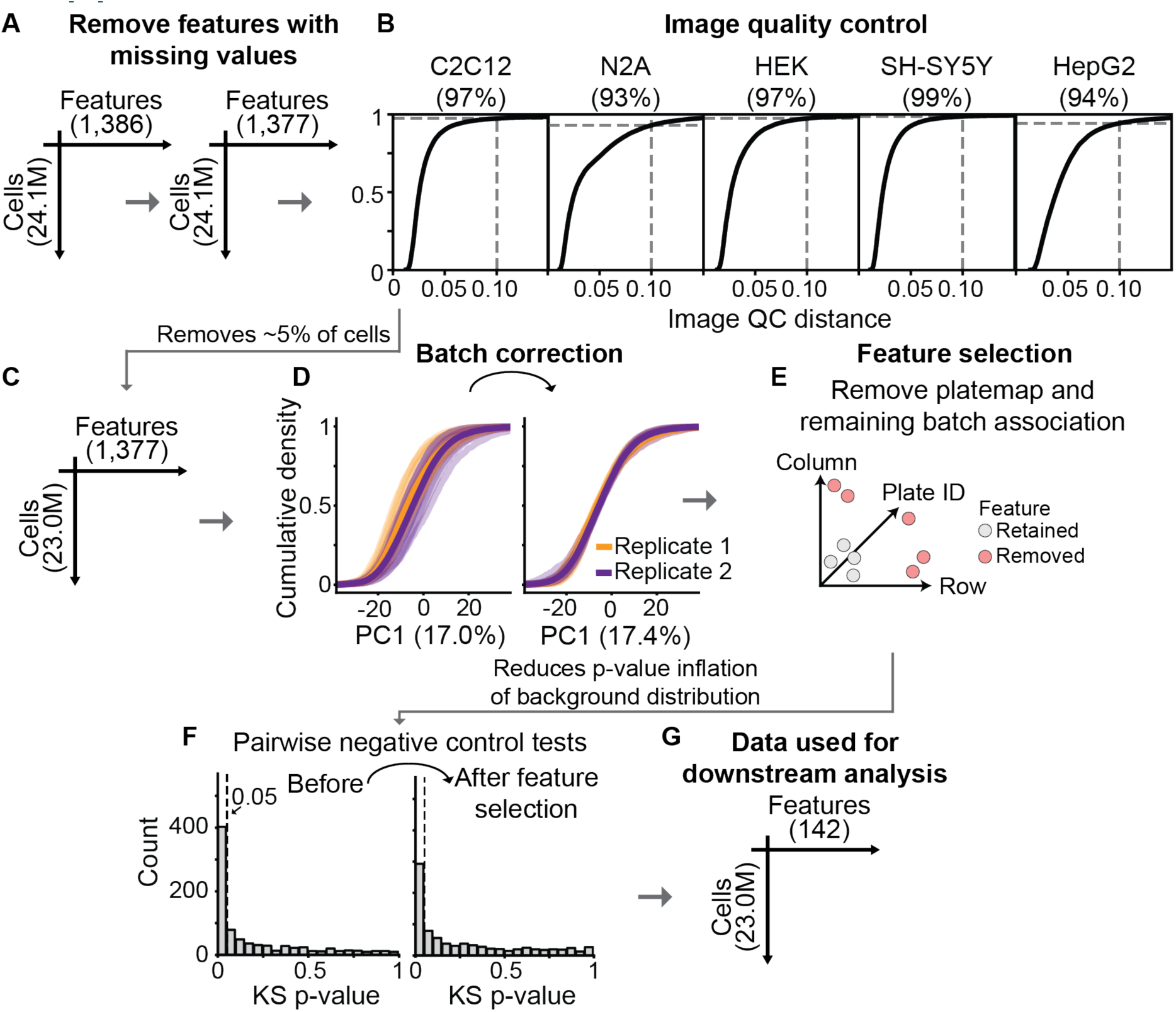
Pre-processing steps of single-cell morphological profiles (see Methods for details). A: Measurements with missing values were removed. B: Image-level features were used for quality control to remove artefacts. Gray lines indicate cutoff beyond which images were discarded due to quality concerns and percentages in title refer to percent of images retained after quality control. C: ∼5% of cells that were in images failing QC were removed, leaving 23M single-cell profiles. D: Batch correction was conducted with a linear model that removed the average between-plate effect per cell line and replicate. E: Kruskal-Wallis statistics were used to identify features affected by plate position or residual batch effect. F: Before step E (feature selection), pairwise tests of negative control wells yielded p-values that are heavily skewed towards zero, indicating false positives. After feature selection, the skew was reduced, although not fully removed. To ensure a well-controlled false discovery rate, we used these p-values as a null distribution (see Methods). G: Together, these steps quality-controlled single-cell morphological profiles, leaving 23M cells and 142 features for analysis (see also Supplementary Table 7).

**Figure S2.**
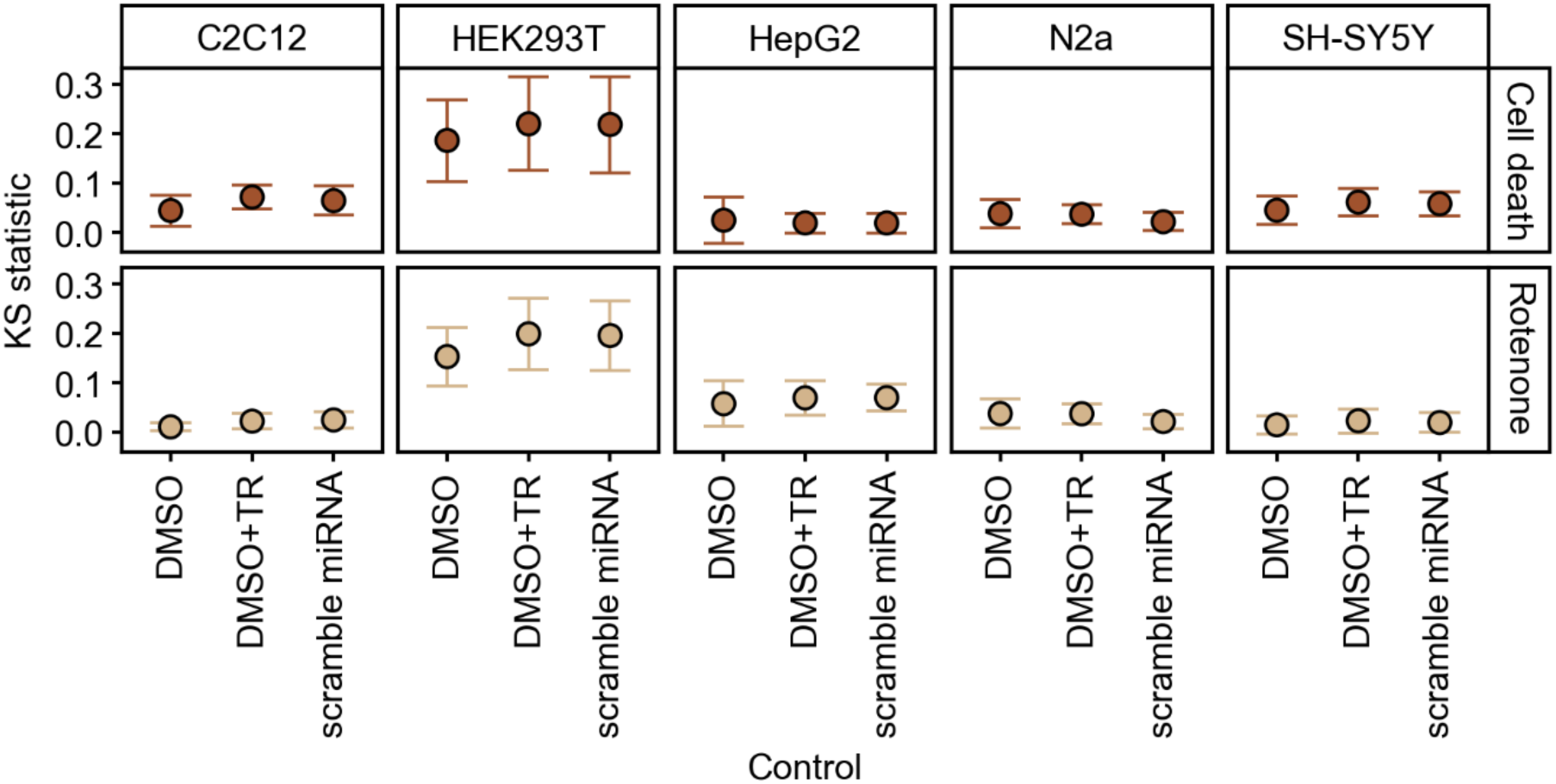
Choice of negative control has only a minor influence on effect size compared to positive controls. Choosing either of the three negative controls as reference and comparing the two positive controls against them shows that the KS statistic, quantifying morphological changes, varies only modestly. Points are mean averages across all plates and experimental replicates; error bars are mean ± standard deviation. Using only DMSO as negative control is associated with slightly larger standard deviation, possibly because there were only four DMSO-only wells per plate, as opposed to eight wells of DMSO with transfection reagent (DMSO+TR) or with additional scrambled miRNA.

**Figure S3.**
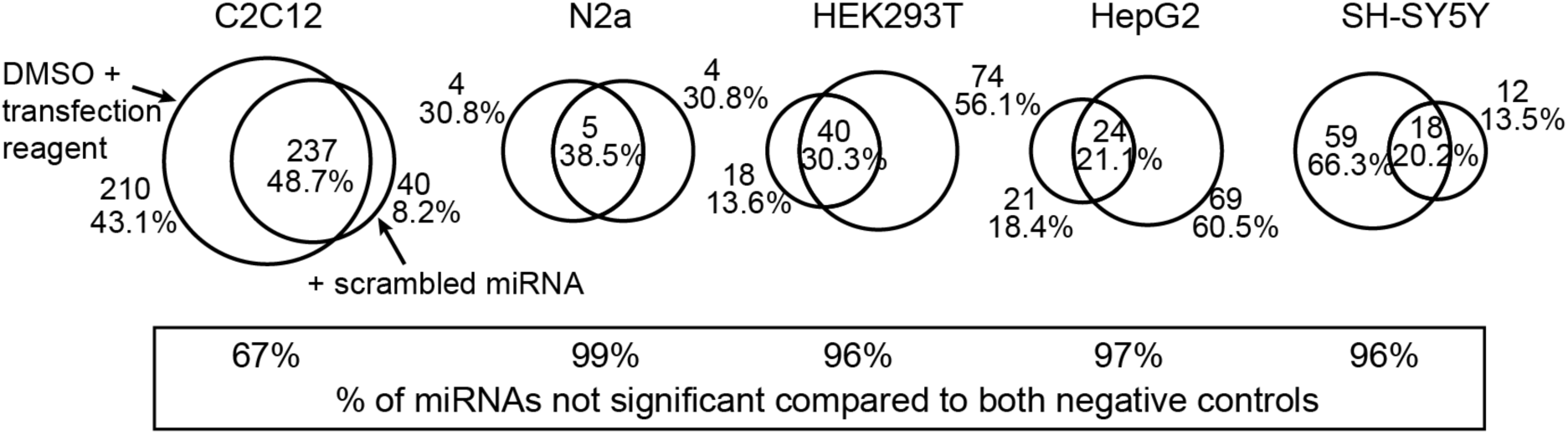
A subset of miRNAs is considered significant compared to two negative controls (intersection) – DMSO + transfection reagent (left circle) and scrambled miRNAs (right circle). In cell lines other than C2C12 there is strong congruence of miRNAs that did not induce significant morphological changes to either negative control (bottom bar).

**Figure S4.**
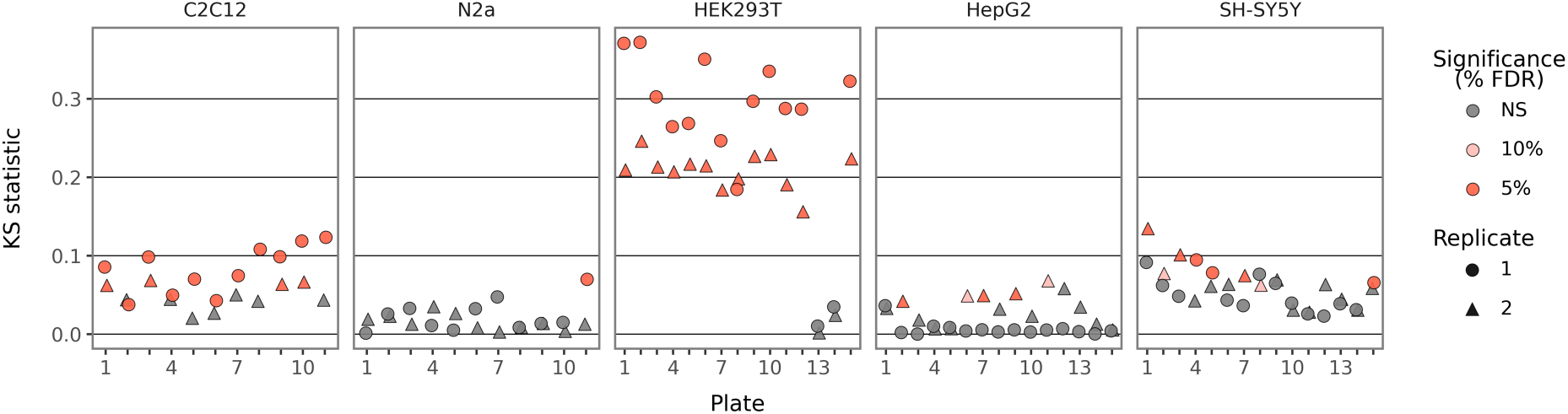
Cell death siRNAs significantly change morphology in HEK293T and C2C12, but less consistently rise above statistical significance in the other cell lines. NS = not significant.

**Figure S5.**
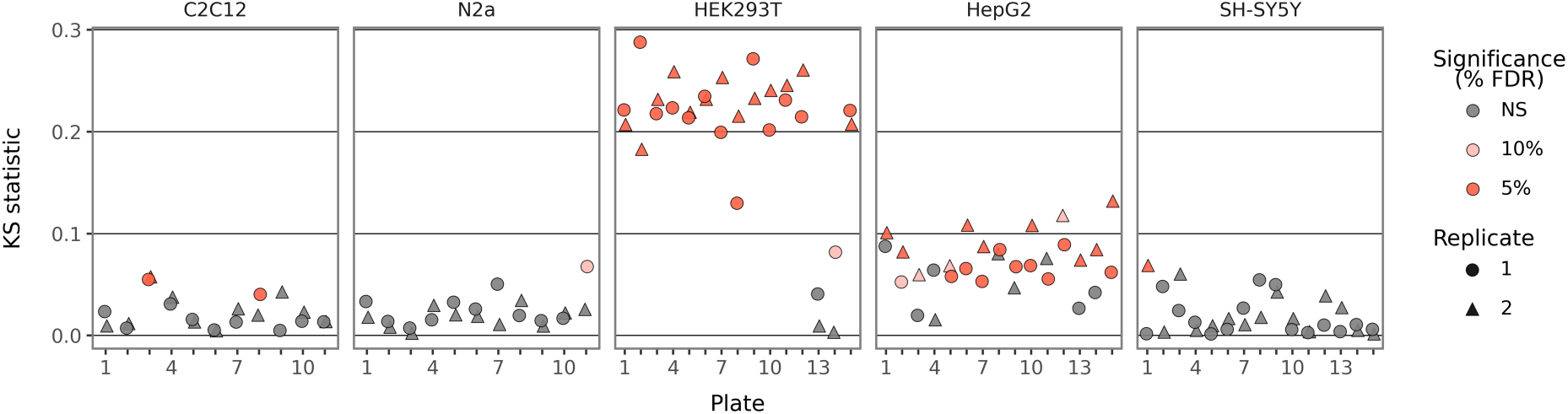
Rotenone significantly changes morphology in most HEK293T and many HepG2 replicates, but not in other cell lines. NS = not significant.

**Figure S6.**
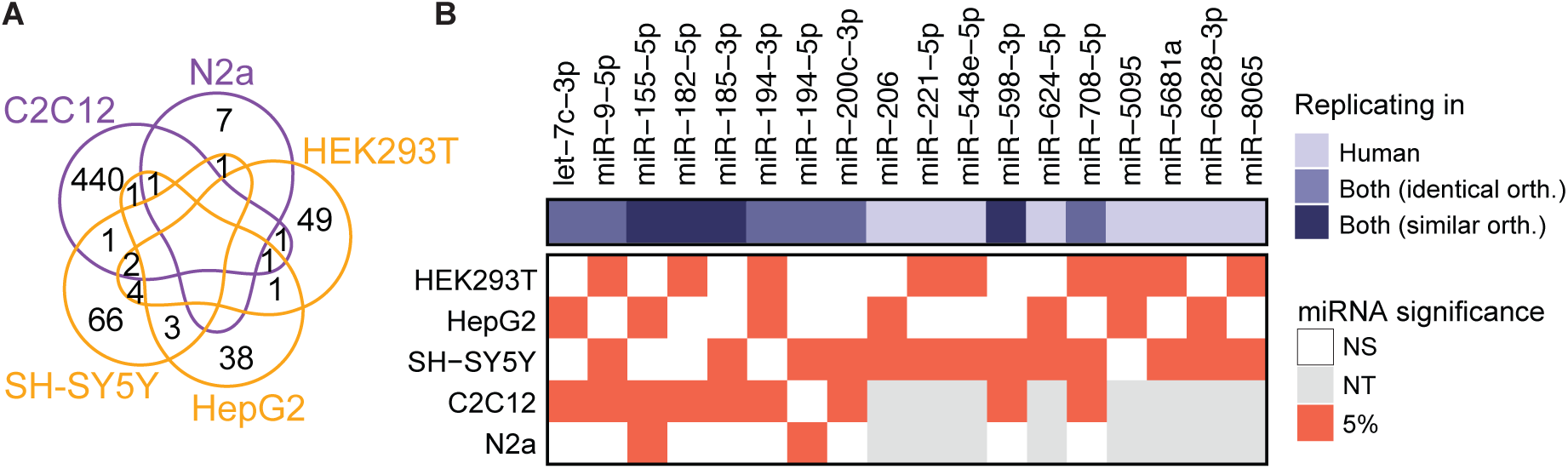
Results for replicating miRNA hits using DMSO + transfection reagent as negative control resemble results shown in Figure 3. A: Venn diagram showing number of miRNAs being a hit in each cell line, including overlaps. Unlabelled segments indicate no shared miRNA. B: 18 miRNAs significantly alter cell morphology in both experimental replicates for each of two cell lines, including six orthologues with perfect sequence identity (“identical orth.”) and three orthologues with sequence identity in the seed and up two 2 nucleotide differences (Levenshtein distance) outside the seed (“similar orth.”). NS = not significant, NT = not tested.

**Figure S7.**
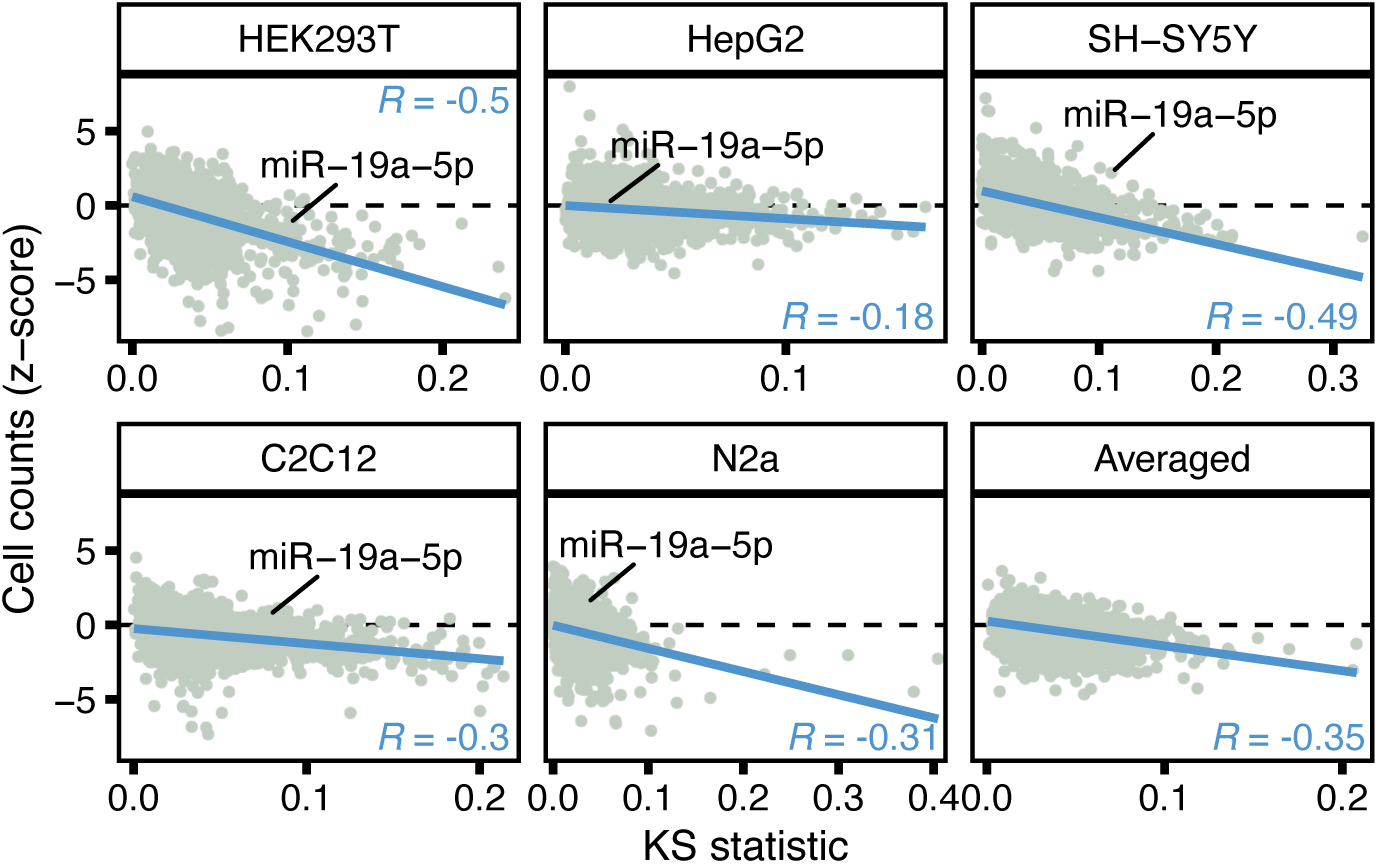
Morphological changes broadly correlate with cytotoxicity, though some miRNAs impact cell morphology without toxicity. Strength of association of morphological changes and cytotoxicity depends on cell line. Each point represents one miRNA; values of the two experimental replicates were averaged. For the “Averaged” panel, data from all cell lines was averaged per treatment. Blue line indicates linear fit.

**Figure S8.**
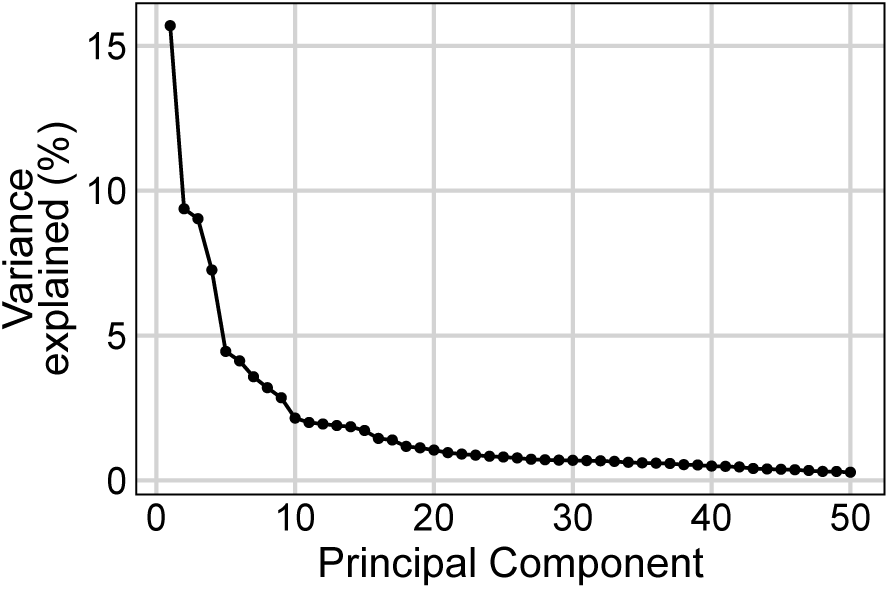
Scree plot for plate 1 of experimental replicate 1 of HEK293T shows that 61.7% of the variance is explained by the first 10 PCs.

## Supplementary Tables

Supplementary Table 1, list of 28 miRNAs removed from miRBase between v21 and v22.1.

Supplementary Table 2, hit calling and cell count results using scrambled miRNA as negative control.

Supplementary Table 3, top 3 differential features by treatment as assessed via t-test using scrambled miRNA as negative control.

Supplementary Table 4, hit calling and cell count results using DMSO+TR as negative control.

Supplementary Table 5, top 3 differential features by treatment as assessed via t-test compared to DMSO+TR as negative control.

Supplementary Table 6, feature names of 155 features retained after feature selection using DMSO+TR as negative control.

Supplementary Table 7, feature names of 142 features retained after feature selection using scrambled miRNA as negative control.

